# Diurnal stomatal apertures profile and density ratios affect whole-canopy conductance, drought response, water-use efficiency and yield

**DOI:** 10.1101/2022.01.06.475121

**Authors:** Sanbon Chaka Gosa, Bogale Abebe Gebeyo, Ravitejas Patil, Ramón Mencia, Menachem Moshelion

## Abstract

Key physiological traits of crop plants, such as transpiration, stomatal conductance, and photosynthesis are highly related to plant productivity. However, these traits are typically studied under steady-state conditions or modeled using only a few measured data points which do not reflect the dynamic behavior of the plant in response to field conditions. In this work, we hypothesized that the plastic behavior of whole-plant water balance regulation, as previously observed in tomato WT, was partially lost in the breeding process due to selective pressure towards productivity. We also hypothesized that this plastic behavior would be observed in some members of the tomato introgression lines (ILs) population, which was created by crossing the WT and M82 lines, particularly in ILs that demonstrate improved drought response. To overcome the steady-state bottleneck, and to test our hypothesis we used a gravimetric functional-phenotyping platform and a reverse-phenotyping method to examine the dynamic whole-plant water-regulation responses of tomato ILs and compared those responses with several years of yield performance in commercial fields. Indeed, our study enabled us to identify high plasticity in a few ideotypic ILs. We found that ideotype lines with highly plastic stomatal conductance and high abaxial-adaxial stomatal density ratios had stomatal apertures that peaked early in the day, even under water-deficit conditions. These traits resulted in dynamic daily water-use efficiency, along with rapid recovery of transpiration when irrigation was resumed after a period of imposed drought. Abaxial stomatal density was also found to be strongly correlated with the expression of the stomatal-development genes *SPCH* and *ZEP*. This study demonstrates how a reverse functional phenotyping approach based on field yield data, continuous and simultaneous whole-plant water-balance measurements and anatomical examination of individual leaves can help us to understand and identify dynamic and complex yield-related physiological traits.

## Introduction

Agricultural crop productivity must increase significantly to meet the food needs of the world’s growing population (FAO, 2017). The challenge of enhancing yields, especially under stressful conditions, while minimizing environmental damage remains paramount (Tian et al., 2021). The most promising strategy to address this is enhancing crop genetics through breeding, informed by genomics and accurate phenotyping of plants and their environmental interactions (i.e., genotype– environment interactions or G×E) [Furbank and Tester (2011), as reviewed in (Gao, 2021)]. Breeding can either narrow genetic diversity by selecting specific germplasm or expand it by introducing alleles from wild relatives. For instance, genetic erosion in tomato has been noted (Schouten et al., 2019). The imprecise quantification of stress-related plant traits has impeded the translation of genomic data into valuable phenotypes (Mir et al., 2019). Thus, the bottleneck in harnessing crop traits for enhanced productivity under stress is transitioning from high-throughput genomics to high-throughput phenomics, which can identify new traits in closely related wild types and predict field yield and resilience.

The first decade of the phenomic era witnessed significant advances in phenotyping platforms. However, the outcomes are not yet optimal, largely due to plants’ highly variable phenotypic responses to their environments (Duursma et al., 2019). Owing to their sessile nature (Claeys and Inzé, 2013), plants are the most adaptable macro-organisms on earth (Schlichting, 1986), showing pronounced phenotypic and physiological adaptability to environmental changes (Moshelion, 2020). Notably, under limited water conditions, physiological adaptability is more vital than morphological adaptability (Marchiori et al., 2017)

Plants display adaptable behavior to complex and unpredictable environmental conditions to optimize their water-use efficiency (WUE) (Hetherington and Woodward, 2003). This adaptability complicates the traditional gene-to-trait research approach since the relationship between genotype and observed traits fluctuates based on environmental conditions (Wardle, 2013). Identifying a gene or QTL contributing to yield, especially drought responses, is challenging due to the intricate nature of yield-related traits. This identification process is time-consuming (Sandhu et al., 2021), limiting the number of promising candidates advancing to field experiments (Moshelion and Altman, 2015). To address these challenges, pre-field functional screening of candidate genotypes can significantly streamline breeding for resistance to abiotic stress (Negin and Moshelion, 2017). These screenings should be brief, mimic field conditions, account for spatial and temporal G×E, and be replicable.

Identifying yield-related traits early in growth is a major challenge in pre-breeding programs (Voss-Fels et al., 2019). Although photosynthesis doesn’t always correlate with yield (Paul, 2021), it often serves as a direct indicator of plant productivity. Thus, the photosynthesis rate is a valuable parameter for predicting yield-related traits and selecting top candidates early in growth (Sallam et al., 2019). For instance, leaf chlorophyll a fluorescence has been a proxy for photosynthesis in various phenotyping platforms (Tietz et al.,2016). However, a reliable high-throughput tool for continuous and simultaneous measurement of whole-plant photosynthetic rates across multiple plants under dynamic conditions is still lacking. As plants regulate productivity-vulnerability trade-offs through stomatal aperture control (Shahinnia et al., 2016), measuring stomatal conductance (gs) and transpiration can be another effective screening strategy. These traits are often studied using steady-state measurements, which don’t capture plant behavior under fluctuating field conditions (Chazdon and Pearcy, 1986; Matthews et al., 2017). A study on rice by Adachi et al., 2019 underscored the influence of dynamic gs on photosynthesis and crop yield, emphasizing the importance of understanding plants’ dynamic responses to their environment.

Factors like stomatal density, size, aperture, distribution, and leaf shape determine levels of gs and transpiration (Ohsumi et al., 2007; Shahinnia et al., 2016). These are challenging to phenotype. Recent studies by Vialet-Chabrand et al., 2013 and Durand et al., 2019 showed that dynamic models predict gs more accurately than steady-state models. Moreover, continuous trait measurements in young tomato plants revealed a close relationship with field yield performance (Gosa et al., 2022).

The dynamics of whole-canopy conductance (Gsc) and its environmental interactions remain elusive. This highlights the need for precise physiological phenotyping for enhanced crop breeding (Ghanem et al., 2015). Whole-plant G×E functional phenotyping platforms are pivotal in addressing the phenomic era’s challenges (Dalal et al., 2020). These platforms offer accurate data on the entire plant’s stomatal behavior (Roman et al., 2021). Stomatal responses are influenced by both external conditions and the plant’s internal biochemical-physiological state. We hypothesize that rapid stomatal responses to environmental changes are vital for optimizing whole-plant momentary WUE. We further propose that this dynamic WUE is crucial for maximizing crop yield, especially under the unpredictable conditions of the growing season. Our rationale is based on observing a morning peak in canopy conductance in Solanum penelli, which seems reduced in the M82 cultivar due to selective breeding (Lupo and Moshelion, 2023).

In recent decades, the emergence of reverse genetics has transformed our understanding of genotype-phenotype relationships in organisms (Walpita and Flick, 2005). Operating similarly to reverse genomics, reverse phenomics was introduced about a decade ago. It involves a detailed analysis of valuable traits to understand their mechanisms, enabling researchers to identify and utilize these traits, shedding light on why certain genotypes or lines excel in specific environments (Furbank and Tester, 2011; Thao and Tran, 2016).

In our study, we employed reverse phenotyping for a multi-year field experiment, analyzing yields and plant weights of tomato introgression lines (IL) where S.penelli genetic material is integrated into the domesticated M82. We monitored the physiological performance of these ILs under both wet and dry conditions using a high-throughput phenotyping system. This enabled us to identify 29 IL lines with a range of key traits, from high yield and resilience to low yield and resilience. We phenotyped the key physiological traits of the best and worst-performing lines early in growth using manual measurements of leaf gas exchange, stomatal imprinting, and an automated functional-phenotyping platform. Our study illustrates how introgression breeding can restore the morning peak of stomatal conductance lost in the commercial M82 cultivar. We also showcased the advantages of continuously and simultaneously tracking whole-plant water-balance regulation traits in response to changing environmental conditions, compared to single time-point measurements. This approach allowed us to connect leaf-level data with whole-plant physiological data measured in greenhouses and open-field yield data, examining the traits.

## Materials and Methods

### Field experiments

We studied 30 tomato genotypes, 29 ILs from crossings of Solanum pennellii with M82 (Solanum lycopersicum cv. M82; Table 1), each containing a unique Solanum pennellii chromosome segment (Eshed and Zamir, 1995). Historical yield data from field experiments between 1993 and 2010 were used for seven selected genotypes. Experiments were conducted at the Western Galilee Experimental Station, Akko, Israel, using a randomized block design (Gur and Zamir, 2004) with a planting density of one plant/m^2^. Irrigation managed drought scenarios, as no rainfall occurred (Fridman et al., 2000; Gur and Zamir, 2004).

**Table 1.** Reverse phenotypic classification of tomato introgression lines based on total yield and plant weight under well-irrigated and dry field conditions (as presented in Fig. 2). Relative to M82, lines were classified into three groups based on their fruit weights: high-yielders (HY, >20%), low-yielders (LY, 20%) and medium-yielders (MY, similar). Similarly, lines were classified as high biomass (HB), medium biomass (MB) and small biomass (SB), relative to M82. Lines with all “highs” were considered ideotypic. The drought stress-response phenotype was classified based on terminology suggested by Moshelion (2020). Specifically, under drought conditions a line with a fruit weight that was 20% higher fruit than that of M82 was classified as having a high tolerance (HT), a line with a fruit weight that was 20% lower than that of M82 was classified as having a low tolerance and lines with fruit weights similar to that of M82 were classified as having moderate tolerance (MT). Plants that maintained or increased their biomass under stress were categorized as having a high resilience (HR). Plants that maintained or reached medium levels of shoot biomass were classified as medium resilience (MR). Small plants that maintained their size or larger plants that lost biomass under stress were classified as having low resilience (LR); the former phenotype was also referred to as a survival phenotype. We identified 20 classification groups; seven of which (marked with *) were selected for further physiologic characterization using the functional telemetric platform. All these lines exhibited consistent behavior across the years of study data.

| line | ID Number | Well irrigation treatment |  | Drought treatment |  | Group classification | Abbreviation | Classification Number |  |
| --- | --- | --- | --- | --- | --- | --- | --- | --- | --- |
|  |  | Fruit yield | Shoot weight | Fruit yield | Shoot weight |  |  |  |  |
| M82 | 30 | CONTROL | CONTROL | CONTROL | CONTROL | Control |  |  | * |
| IL12-1-1 | 4 | HIGH | HIGH | HIGH | HIGH | High yielder High biomass High tolerance High resilience - IDEOTYPE | HyHbHtHr | 1 |  |
| IL5-2 | 16 | HIGH | HIGH | HIGH | HIGH | High yielder High biomass High tolerance High resilience - IDEOTYPE | HyHbHtHr | 1 | * |
| IL6-4 | 26 | HIGH | HIGH | HIGH | HIGH | High yielder High biomass High tolerance High resilience - IDEOTYPE | HyHbHtHr | 1 |  |
| IL7-4-1 | 22 | HIGH | HIGH | HIGH | HIGH | High yielder High biomass High tolerance High resilience - IDEOTYPE | HyHbHtHr | 1 |  |
| IL2-4 | 8 | HIGH | HIGH | MEDIUM | HIGH | High yielder High biomass Medium tolerance High resilience | HyHbMtHr | 2 |  |
| IL2-6 | 9 | HIGH | HIGH | LOW | HIGH | High yielder High biomass Low tolerance High resilience | HyHbLtHr | 3 | * |
| IL11-4 | 3 | HIGH | MEDIUM | MEDIUM | MEDIUM | High yielder Medium biomass Medium tolerance Medium resilience | HyMbMtMr | 4 | * |
| IL6-3 | 20 | MEDIUM | HIGH | LOW | HIGH | Medium yielder High biomass Low tolerance High resilience | MyHbLtHr | 5 |  |
| IL9-3 | 29 | MEDIUM | HIGH | LOW | HIGH | Medium yielder High biomass Low tolerance High resilience | MyHbLtHr | 5 |  |
| IL9-1 | 26 | MEDIUM | HIGH | LOW | MEDIUM | Medium yielder High biomass Low tolerance Medium resilience | MyHbLtMr | 6 |  |
| IL9-2-6 | 28 | MEDIUM | HIGH | MEDIUM | HIGH | Medium yielder High biomass Medium tolerance High resilience | MyHbMtHr | 7 |  |
| IL2-6-5 | 10 | MEDIUM | MEDIUM | HIGH | HIGH | Medium yielder Medium biomass High tolerance High resilience | MyMbHtHr | 8 |  |
| IL5-3 | 17 | MEDIUM | MEDIUM | MEDIUM | HIGH | Medium yielder Medium biomass Medium tolerance High resilience | MyMbMtHr | 9 |  |
| IL9-1-3 | 27 | MEDIUM | MEDIUM | MEDIUM | LOW | Medium yielder Medium biomass Medium tolerance Low resilience | MyMbMtLr | 10 |  |
| IL10-2 | 2 | MEDIUM | MEDIUM | MEDIUM | MEDIUM | Medium yielder Medium biomass Medium tolerance Medium resilience | MyMbMtMr | 11 |  |
| IL5-5 | 18 | MEDIUM | SMALL | HIGH | HIGH | Medium yielder Small biomass High tolerance High resilience | MySbHtHr | 12 |  |
| IL2-1-1 | 7 | MEDIUM | SMALL | MEDIUM | LOW | Medium yielder Small biomass Medium tolerance Low resilience | MySbMtLr | 13 |  |
| IL12-2 | 5 | MEDIUM | SMALL | MEDIUM | MEDIUM | Medium yielder Small biomass Medium tolerance Medium resilience | MySbMtMr | 14 |  |
| IL12-3-1 | 6 | MEDIUM | SMALL | MEDIUM | MEDIUM | Medium yielder Small biomass Medium tolerance Medium resilience | MySbMtMr | 14 |  |
| IL10-1 | 1 | LOW | HIGH | LOW | HIGH | Low yielder High biomass Low tolerance High resilience | LyHbLtHr | 15 | * |
| IL3-3 | 11 | LOW | HIGH | LOW | HIGH | Low yielder High biomass Low tolerance High resilience | LyHbLtHr | 15 |  |
| IL3-4 | 12 | LOW | HIGH | LOW | HIGH | Low yielder High biomass Low tolerance High resilience | LyHbLtHr | 15 |  |
| IL6-2 | 19 | LOW | MEDIUM | LOW | LOW | Low yielder Medium biomass Low tolerance Low resilience | LyMbLtLr | 16 |  |
| IL4-1 | 13 | LOW | MEDIUM | MEDIUM | MEDIUM | Low yielder Medium biomass Medium tolerance Medium resilience | LyMbMtMr | 17 |  |
| IL4-1-1 | 14 | LOW | SMALL | MEDIUM | MEDIUM | Low yielder Small biomass Medium tolerance Medium resilience | LySbMtMr | 18 |  |
| IL8-1 | 23 | LOW | SMALL | LOW | MEDIUM | Low yielder Small biomass Low tolerance Medium resilience | LySbLtMr | 19 | * |
| IL8-1-3 | 25 | LOW | SMALL | LOW | LOW | Low yielder Small biomass Low tolerance Low resilience- SURVIVAL | LySbLtLr | 20 | * |
| IL4-3-2 | 15 | LOW | SMALL | LOW | LOW | Low yielder Small biomass Low tolerance Low resilience - SURVIVAL | LySbLtLr | 20 |  |
| IL8-1-1 | 24 | LOW | SMALL | LOW | LOW | Low yielder Small biomass Low tolerance Low resilience- SURVIVAL | LySbLtLr | 20 |  |

### Leaf gas-exchange measurements

Plants were grown in a semi-controlled greenhouse at the Hebrew University of Jerusalem. Leaf gas-exchange was assessed on the youngest, fully extended leaf of ∼8-week-old plants using a portable infrared gas analyzer (LI-6800XT; Li-Cor Inc.). Conditions mirrored the greenhouse’s environment.

### Reverse phenomics in a greenhouse

#### Experimental setup

Eight selected ILs and M82 tomato plants were grown in a greenhouse at The Hebrew University of Jerusalem in September 2019. Details on nutrients and setup are in (Dalal et al., 2020). Highly sensitive load cells, part of the Plantarray system (PlantDitech, Yavne, Israel), were used as weighing lysimeters, each supporting a 4-L pot with a single plant. Fertilizer was supplied via fertigation.

#### Drought treatment

To ensure uniform drought treatment, the system irrigated each plant to 80% of its previous day’s transpiration [see Figure 3B in (Dalal et al., 2020)].

#### Measurement of quantitative physiological traits

Plant water-relations kinetics and physiological traits were determined for all plants (Figure 1), following (Halperin et al., 2017) with modifications. Traits included daily transpiration, transpiration rate (TR), whole-canopy stomatal conductance (Gsc), and biomass water-use efficiency (WUEb). Recovery from drought was assessed by comparing daily transpiration post-recovery.

**Figure 1.**
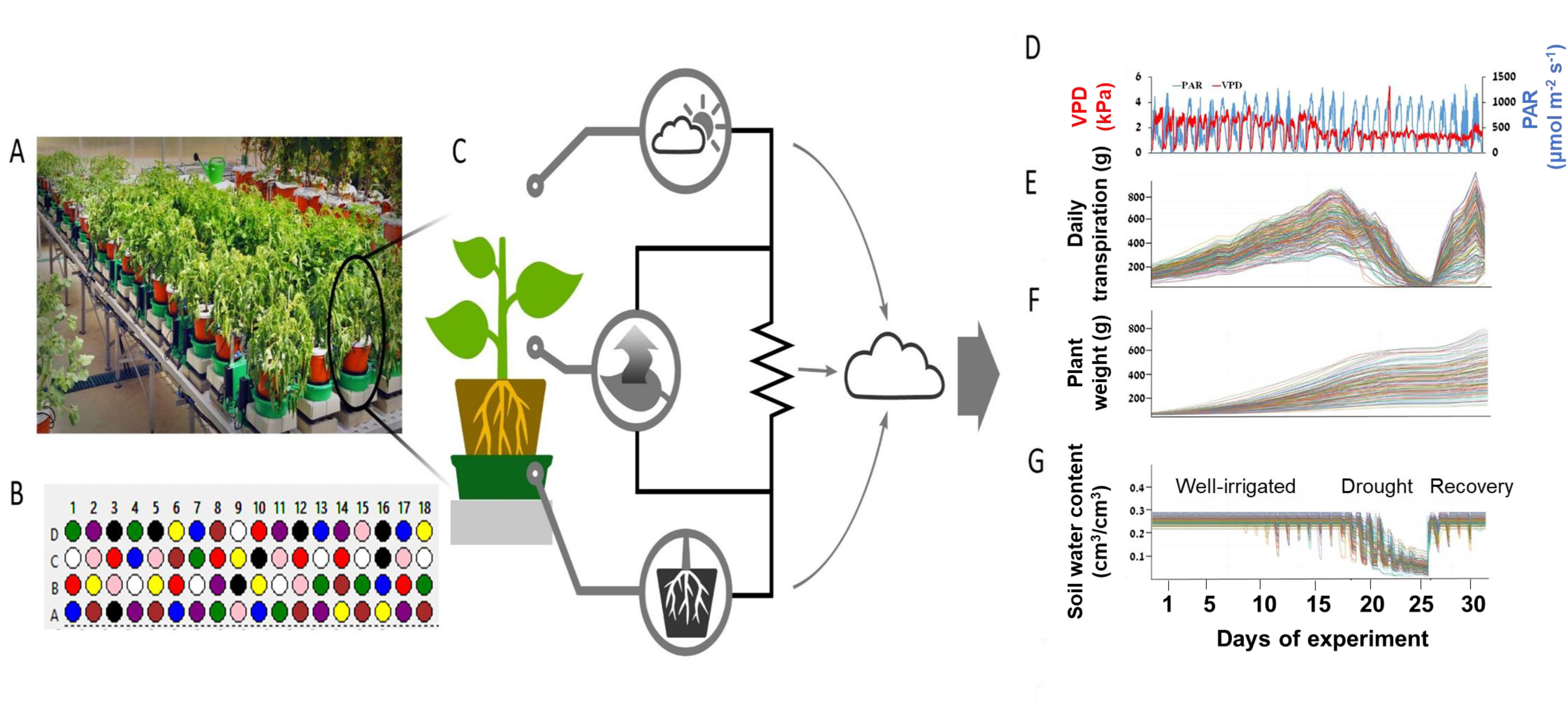
Overview of the telemetric, gravimetric phenotyping platform and analysis scheme. A, A partial view of multiple tomato introgression lines positioned on the Plantarray screening platform [located at the Israeli Center of Research Excellence (ICORE) for Plant Adaptation to the Changing Environment, at The Hebrew University of Jerusalem]. B, Randomized experimental setup of all plants simultaneously measured. Different colors represent different lines and treatments. C, An illustration of the direct soil-plant-atmosphere measurements taken for a single plant. The continuous data from the Plantarray system is uploaded to the internet server in real time. D, Graphic presentation of the absolute values of the continuous atmospheric data of vapor pressure deficit (VPD, red line) and photosynthetically active radiation (PAR, blue line) over time. E, Whole-plant daily transpiration kinetics. F, Plant weight gain over time G, soil moisture content over time. Each line represents the continuous measurement of individual plants throughout the experiment period.

#### Stomatal density and aperture

Stomatal apertures and density were determined using a rapid imprinting method (Geisler and Sack, 2002) (Figures 8 and 9). Imprints were analyzed using ImageJ software with size calibration via a microscopic ruler.

#### Statistical analysis

Data were processed using SPAC analytic software (PlantDitech, Yavne, Israel). Analyses were performed using JMP® 15.0 Pro (SAS Institute). Graphs were generated with OriginPro, Version 2021 (OriginLab Corporation).

**For a full description, see supplemental materials and methods.**

## Results

### Field performance of the IL population

We began by analyzing two field experiments (from 2000 and 2004, see Table 1) that included 29 IL lines and M82. These plants were characterized for yield and biomass parameters and compared to the control M82 under optimal irrigation and water-limiting conditions (Figure 2). Under optimal irrigation, the lines were classified into high-yielder (HY), medium-yielder (MY), and low-yielder (LY) groups based on their yields relative to M82: HY = >20% of M82 yield, MY = like M82 yield, and LY = <20% of M82 yield. Based on the biomass of the different lines collected from the field at the experiment’s end relative to M82, the genotypes were also classified as having high shoot biomass (HB, >20%), medium shoot biomass (MB, 20%), or low shoot biomass (LB, <20%; Table 1).

**Figure 2.**
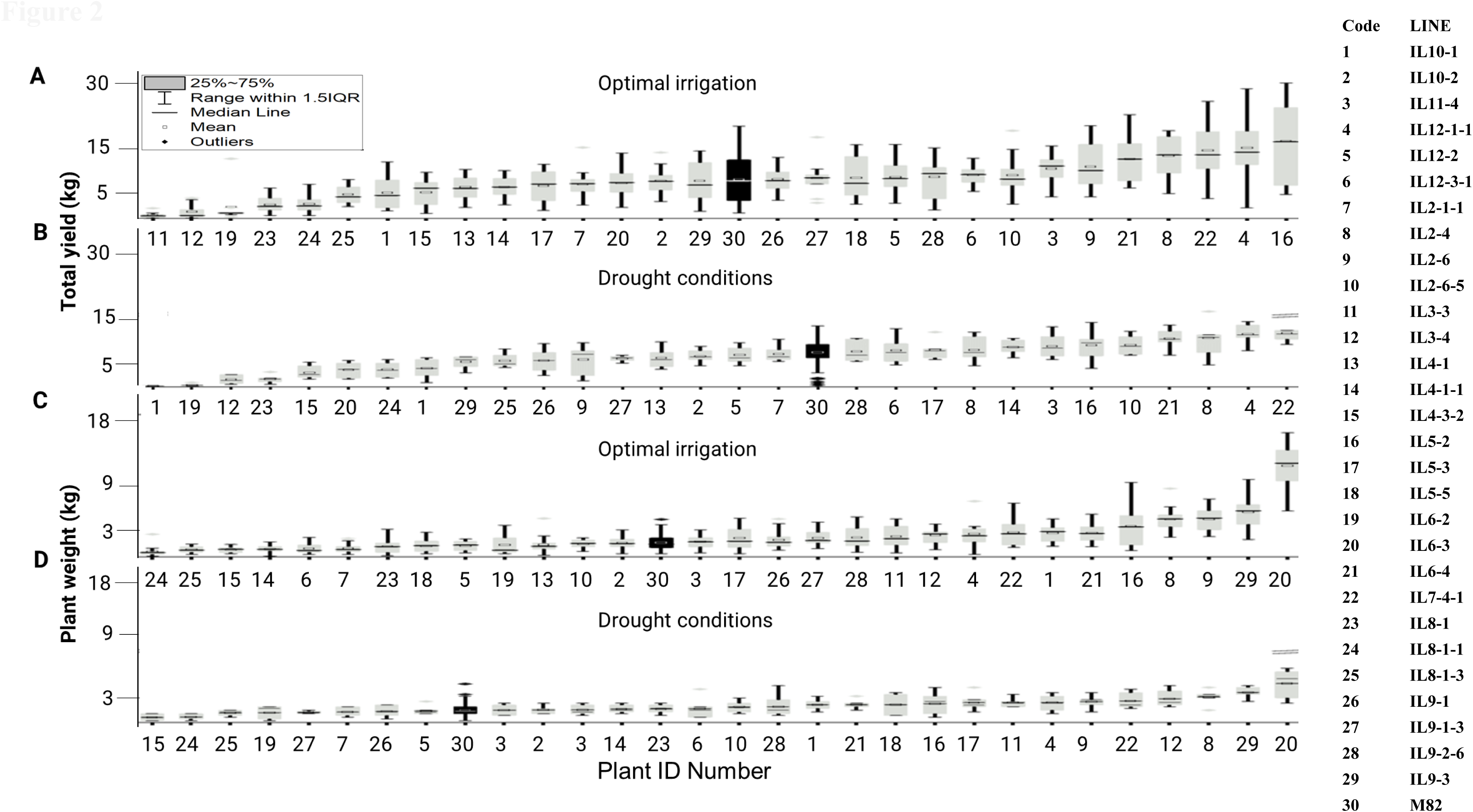
Field performance of 29 tomato introgression lines (gray) and the M82 control (black). A, Two years of mean fruit weights (ranked from low to high) under well-irrigated conditions. B, Two years of mean fruit weights (ranked from low to high) under drought conditions. C, Two years of mean shoot weights at harvest (ranked from low to high) under well-irrigated conditions. D, Two years of mean shoot weights at harvest (ranked from low to high) under drought conditions. The square (□) in the box plot represents the mean value. The box-splitting horizontal bands indicate the sample median and bars show the interquartile range (25th to 75th percentile). Points below or above the interquartile ranges are outliers respectively.

Following the classification and the terminology proposed by (Moshelion, 2020), plant drought-response behavior was defined as follows: Plant resilience was gauged by the plant’s biomass relative to M82, a check variety under similar drought conditions, and plant tolerance was measured by its total yield (TY) under similar drought conditions. Thus, under comparable drought stress, a line displaying higher biomass and higher yield would be classified as having a high-resilience (HR) and high-tolerance (HT) phenotype (HrHt). In contrast, a line with lower biomass and yield under drought conditions would be classified as having a low-resilience (LR) and low-tolerance (LT) phenotype (LrLt). If a line’s biomass under stress was >20% of the control, but its yield was lower than M82, it was defined as high-resilience– low-tolerance (HrLt; Table 1). This categorization resulted in 20 different groups, ranging from an ideotypic phenotype to a survival phenotype.

Six lines (marked with ‘*’ in Table 1) and the M82 line were chosen for further physiological characterization using the functional telemetric platform. These seven genotypes were selected to represent a broad range of plant-response characteristics based on data from 4 to 5 years of well-irrigated field experiments and at least two years of data on performance under stress conditions. The selected lines were IL5-2, IL2-6, IL11-4, IL10-1, IL8-1, and IL8-1-3. IL5-2 outperformed M82 in all parameters in both environments. It was an HY, HB, HT, and HR (HyHbHtHr) line, making it ideotypic. IL2-6 was a high yielder under optimal irrigation but a low-yielder under drought stress, while maintaining high biomass under both conditions (HyHbLtHr). IL11-4 exhibited high yields under optimal irrigation but had medium-level values for the other selection criteria (HyMbMtMr). IL10-1 demonstrated robust vegetative growth but had low yields in both wet and dry conditions. It was a low yielder with high biomass, low tolerance, and high resilience (LyHbLtHr). IL8-1 had low total yields and biomass under optimal irrigation and low yields under drought conditions, but medium biomass under drought stress (LySbLtMr). IL8-1-3 had low total yields and biomass under optimal irrigation and low yields under both optimal irrigation and drought conditions (LySbLtLr). We refer to its phenotype as a survival phenotype.

### Physiological characterization of the selected genotypes

The midday A_N_ and gs of 8-week-old plants of selected genotypes grown in sandy-loam soil in a semi-controlled greenhouse revealed no significant differences between these genotypes (Supplemental Figure 1). There was considerable variability among the results, potentially due to spatial and temporal differences in ambient conditions and the plant’s physiological status between measurements. Consequently, we decided to measure all genotypes’ water relations simultaneously and continuously.

All lines, with four to eight biological replications, displayed linear increases in transpiration over the period of optimal irrigation. On the experiment’s first day (4-week-old seedlings), no significant differences were observed between the lines (Figure 3 A, Day 1). However, differences between the lines soon emerged. As the plants grew, some lines, such as IL11-4 and IL5-2, exhibited greater whole-plant transpiration, while others, like IL8-1, showed medium to low levels of transpiration. After 18 days, all plants underwent controlled drought stress (a differential-feedback-irrigation drought treatment, see Materials and Methods), ensuring they all experienced a similar drought-stress treatment wherein soil volumetric water content (VWC) decreased at a similar rate (Figure 3 B), despite their transpiration differences. Although all plants encountered a similar declining VWC, high-transpiring lines reached their VWC limitation point (θcrit.) more swiftly, thus reducing their transpiration earlier than the low-transpiring plants (Figure 4, Suppl. Figure 3), revealing a transpiration-positional inversion (e.g., Figure 3 A, the transpiration flip-flop of IL11-4 and IL8-1).

**Figure 3.**
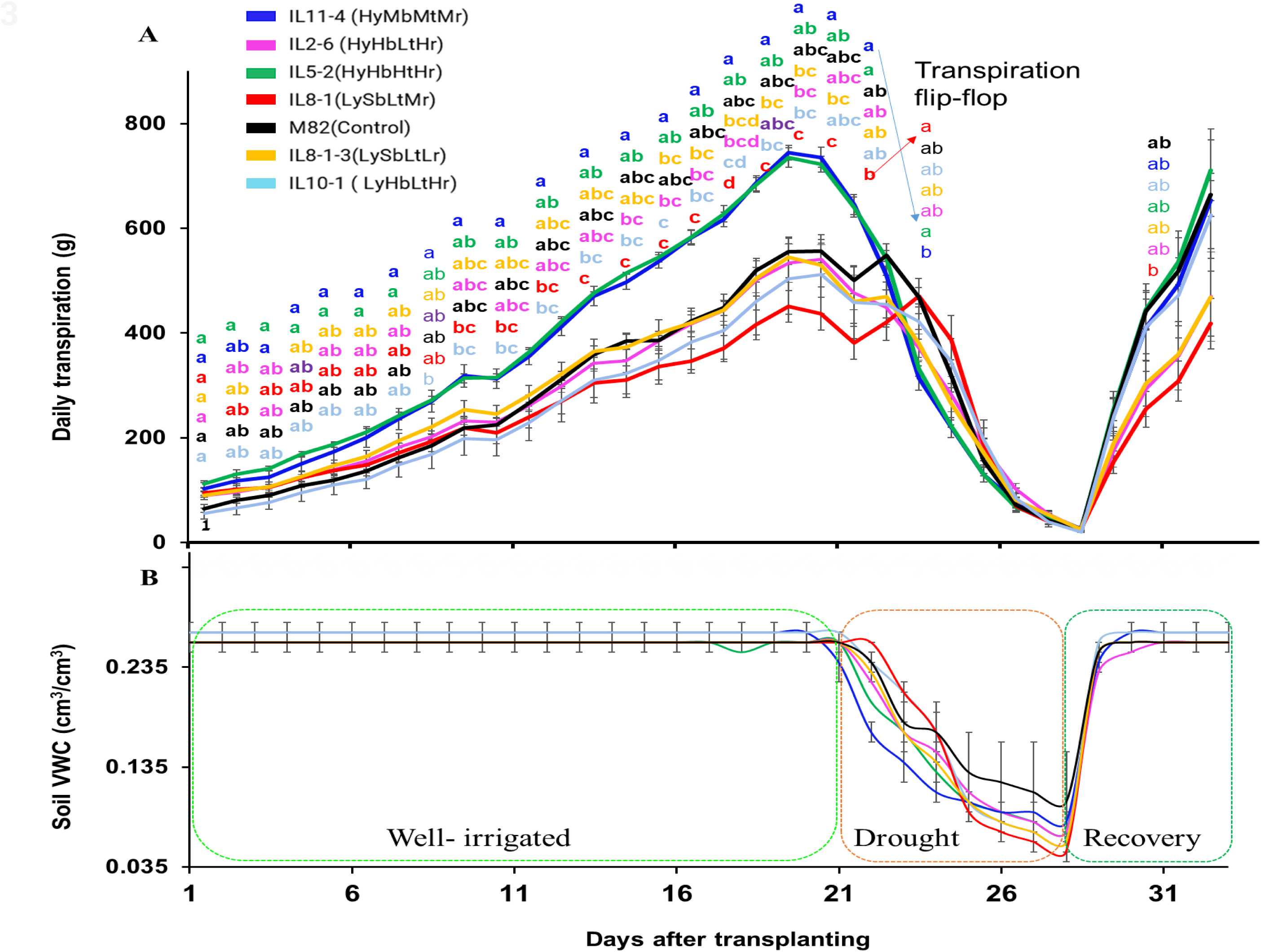
A, Daily transpiration of tomato seedlings over the entire experimental period. Data points are means ± SE of continuous daily whole-plant transpiration over the entire experimental period (32 days). B, Mean ± SE volumetric water content (VWC) measured by a soil probe over the course of the experiment. The drought treatment was followed by a recovery period during which the resumed irrigation brought the pot back to full capacity. Groups were compared using Tukey’s Honest Significance test; different letters above points represent significant differences between lines; *p* < 0.05. *n* = 5–8 plants per group.

**Figure 4.**
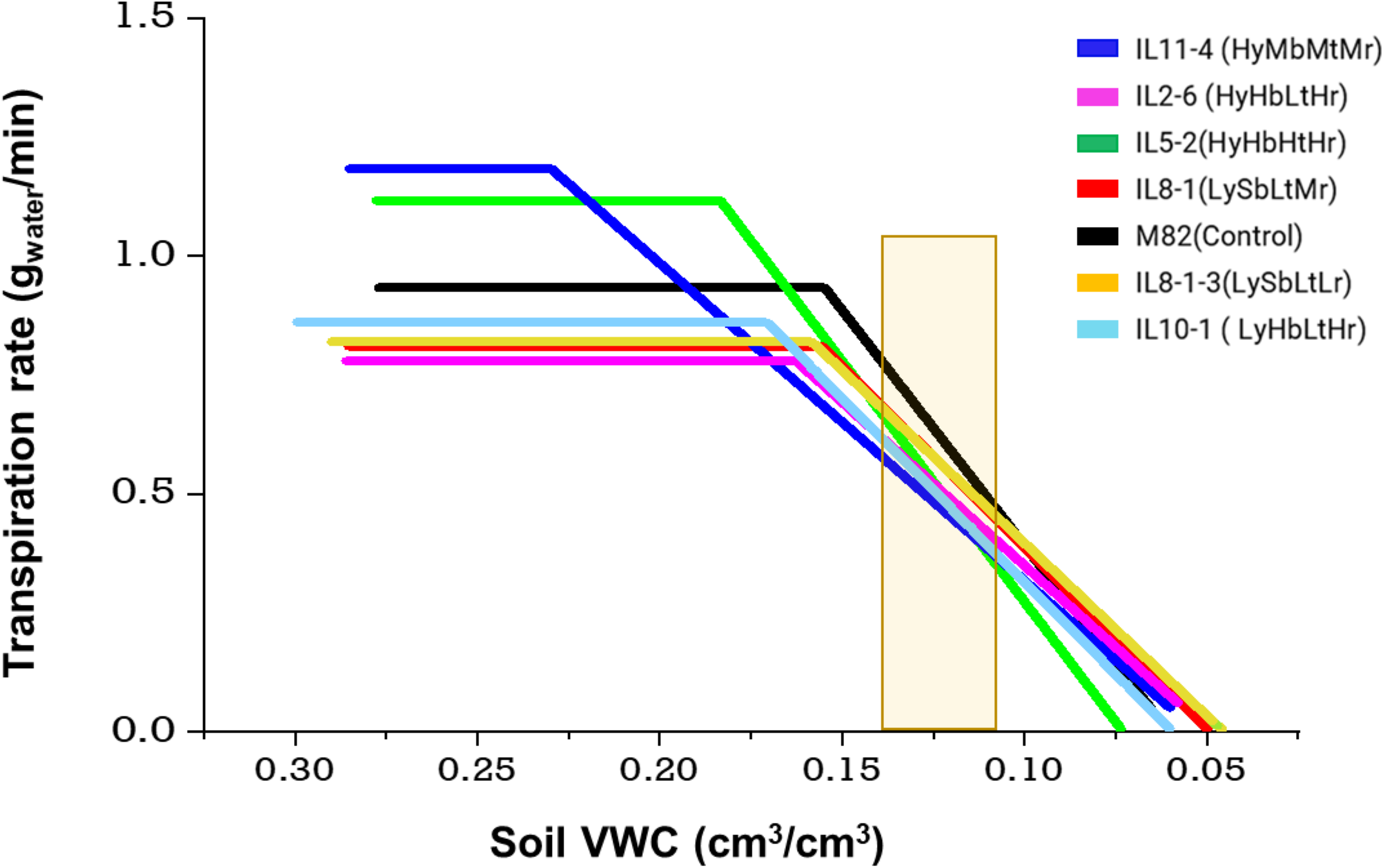
The physiological drought point (θcrit.) was determined as the point at which soil water is restricted from supplying mid-day transpiration needs, for all of the plants presented in Fig. 3. This point was identified using piecewise correlations based on the relationships between two segmented lines that intersected (following Halperin et al., 2016). The yellow box in the middle represents the standard drought-evaluation zone, in which the performance of all lines was evaluated under drought stress.

### Stress response and resilience evaluation

During the drought period, no differences in plant transpiration were observed between the lines (Figure 3 A) due to stomatal closure. It’s highly probable that other drought-defense mechanisms, such as reactive oxygen species (ROS) scavenging, embolism repair, and prevention, were also activated. However, estimating the effectiveness of these mechanisms across all plants in real-time is challenging in intact plants. Thus, we tested the recovery rates of the different lines, assuming that the more effective these mechanisms, the faster the plants would recover. When the plants reduced their transpiration to 10% of their maximum transpiration (Day 29), full irrigation resumed, and the plants’ transpiration was monitored for 5 days. Over this period, IL5-2, IL11-4, IL10-1, and M82 exhibited rapid recovery (Figures 3A, 5A).

The strong correlation between cumulative transpiration (CT) and the amount of dry biomass at the experiment’s end (an integral of the whole-plant transpiration kinetics over the entire 33-day period) revealed that the higher-biomass lines also transpired more than the smaller plants. We observed a linear relationship between cumulative transpiration and total biomass production (Figure 5 B). Moreover, the lines with higher CT (IL5-2 and IL11-4) were also more efficient, as they had higher water-loss to biomass-gain ratios (biomass water use efficiency, WUEb) than the other lines (Figure 5 C).

**Figure 5.**
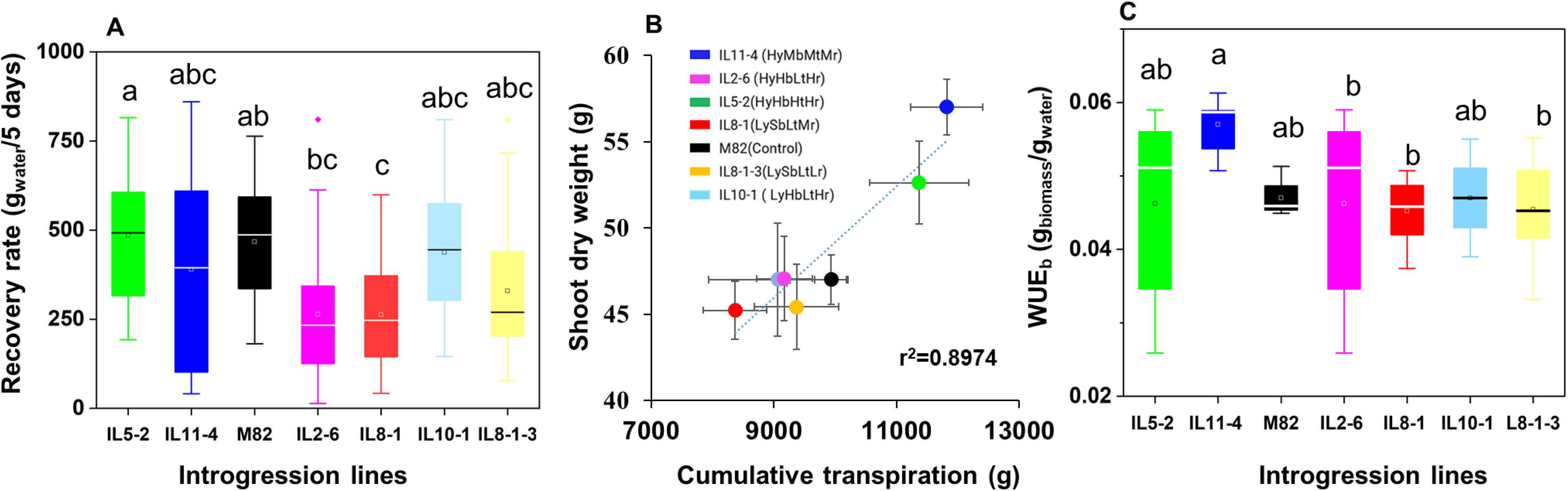
A, Transpiration recovery rate for five days after the resumption of irrigation in seven different tomato lines. B, Correlation between shoot dry weight and cumulative daily transpiration for 33 days during the screening period. Data points are the means ± SE (*n* = 5‒8). C, means ± SE of biomass WUE of the different lines (calculated by dividing the dry biomass weight by the cumulative transpiration of each plant).

### Daily whole-plant–environment kinetics

Despite the qualitative increases in daily transpiration, discerning a clear statistical difference between the lines based on their daily transpiration parameters was challenging. We hypothesized that differences in the momentary plasticity response to the ambient conditions (which were similar for all lines) throughout the day might explain the performance differences between lines. To comprehend the response of Gsc and the transpiration rate to ambient conditions, we monitored all lines simultaneously under optimal irrigation and drought conditions (50% of maximum midday transpiration, after θcrit., Day 23 of the experiment, Figures 6 and 7). Under well-irrigated conditions (Day 16 of the experiment), the whole-canopy Gsc and TR displayed a similar response pattern to light and VPD across different lines. All lines opened their stomata in response to PAR and VPD, increasing their Gsc and TR from 06:00 to 10:00 (Figure 6). Daily maximum Gsc and RT remained relatively stable at midday (10:00 to 14:00) and until late afternoon, consistent with the PAR and VPD. In the late afternoon, Gsc sharply declined in response to PAR, while TR increased following the VPD pattern. With the VPD drop, transpiration also decreased (Figure 6C). This indicates that during the afternoon and evening hours, the plants’ WUE was at its lowest level. The transpiration rate had a pattern like that of the daily transpiration shown in Figure 3 A, but canopy conductance had a different pattern since it was normalized to plant weight. Unlike the situation under optimal-irrigation conditions, the Gsc and TR kinetics of the plants exposed to soil water-limiting conditions varied between the different lines and in response to PAR and VPD (which were very similar to the pre-stress conditions, see Supplemental Figure 2). All lines experienced reductions in Gsc and TR from late morning-noon, regardless of the ambient PAR and VPD conditions. However, the high-performing lines (i.e., IL5-2 and IL11-4) presented a different response pattern, with relatively high Gsc (Figure 7 A and B) in the morning (6:00 to 10:00) and a relatively low transpiration rate (Figures 7 a and b) at the same time, followed by an immediate and linear reduction in those parameters during the middle of the day and the afternoon (10:00 to the evening hours), suggesting a possibly higher WUE for this period compared to other lines. Conversely, low-performing lines (i.e., IL10-1, IL8-1, and IL8-1-3) presented lower Gsc during the early hours (when VPD is low) and reached their Gsc peaks later in the day (Figure 7 E, F, and G, respectively), and those peaks persisted for longer periods, causing those plants to lose more water to transpiration (Figure 7 e, f, and g respectively). In these high-resolution measurements of Gsc and Tr patterns, we detected differences between lines that were not evident in the low-resolution measurements of daily transpiration (i.e., no differences between lines were detected in the daily-transpiration stress response unlike the pre-stress period; Figure 3 A). To better understand the different Gsc response patterns of the high-yielding lines, we examined their stomatal densities and apertures.

**Figure 6.**
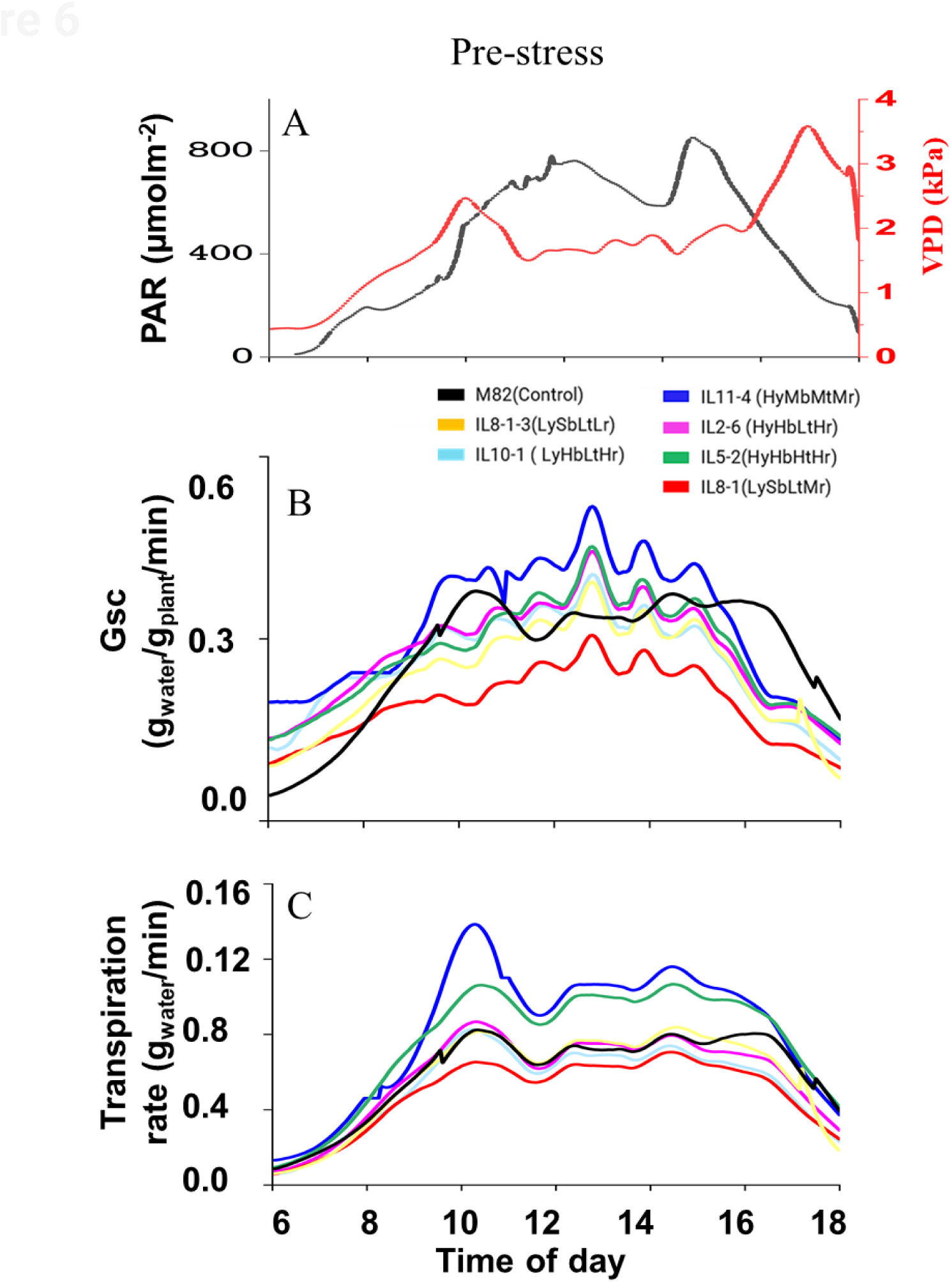
Daily patterns of A, PAR (black line) and VPD (red line), B, whole-canopy conductance and C, whole-canopy transpiration rate, as continuously measured under well-irrigated conditions. Data means ± SE; *n* = 5–8.

**Figure 7.**
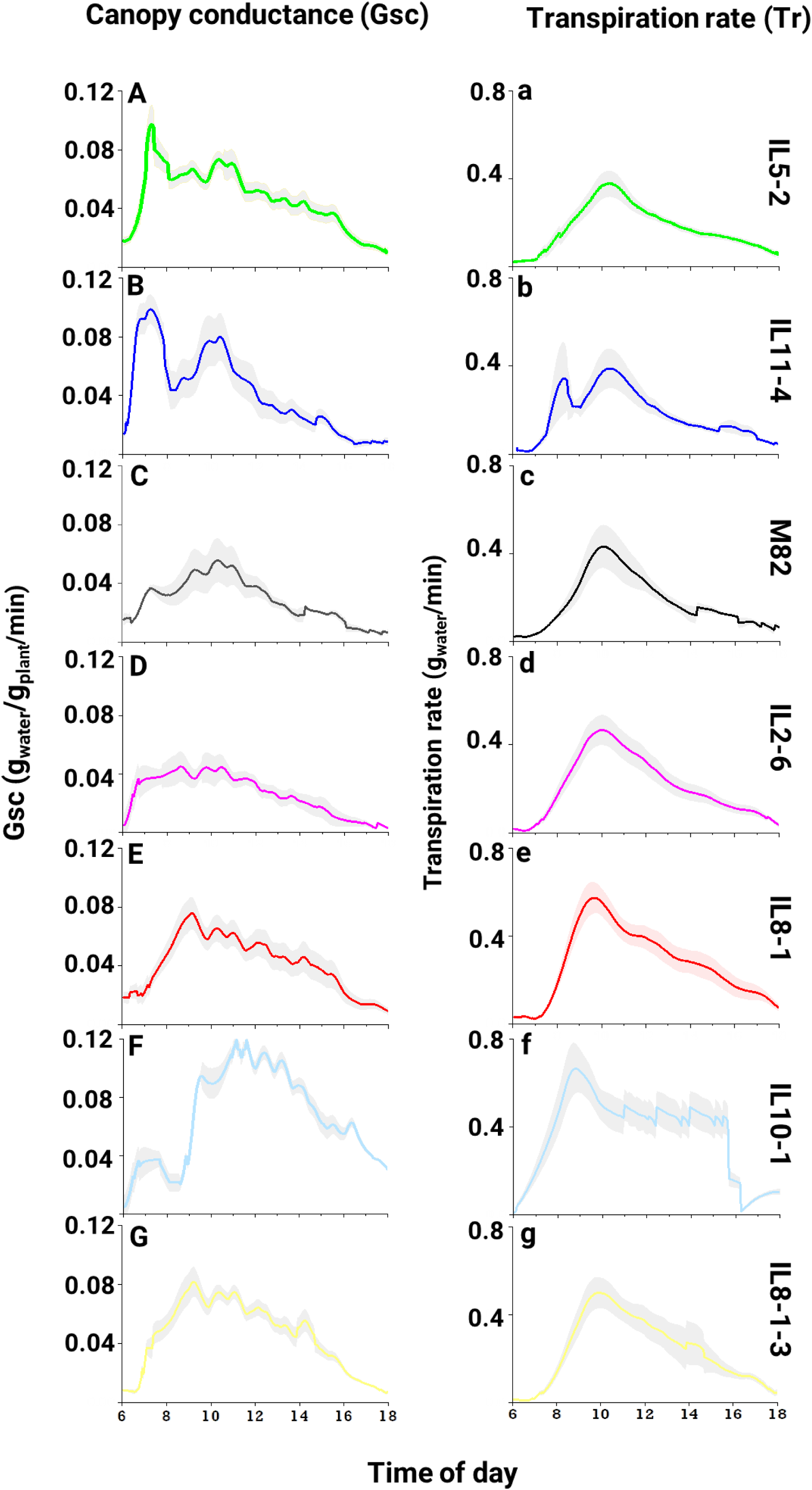
Daily variation in canopy conductance and transpiration rate under drought-stress conditions. Whole-canopy conductance (A–G) and transpiration rate (a–g) measured continuously. Representative days were shown for the stress after the θcrit. point (yellow block in Fig. 4). Data are shown as means ± SE (*n* = 3–8).

### Variation in stomatal morphology among the lines

Stomatal density and aperture measurements revealed that lines IL5-2, IL11-4, IL8-1-3, and IL10-1 had higher abaxial stomatal densities than adaxial stomatal densities. In contrast, the other lines had similar stomatal densities on both leaf sides. No differences were observed in the maximal stomatal aperture size between the abaxial and adaxial leaf sides for each line, except for M82 and IL8-1-3, which had larger stomatal apertures on the abaxial and adaxial sides of their leaves, respectively (Figure 8 B). However, each line had its maximum aperture for both abaxial and adaxial sides of its leaves at different times of day (Figure 9). Specifically, IL5-2 reached its maximal abaxial peak aperture around 07:00 and its maximal adaxial peak aperture around 10:00 (Figure 9 A, a). M82 reached both its maximal abaxial and adaxial apertures around 10:00. Lines IL8-1, IL11-4, and IL10-1 revealed maximum aperture on both their adaxial and abaxial sides around 13:00. However, while the stomatal apertures of IL10-1 and IL11-4 shrank after peaking, IL8-1 maintained its large apertures until 17:00. Line 8-1-3 consistently maintained small apertures throughout the day, with slightly larger apertures in the afternoon on the adaxial side of its leaves (Figure 9 G and g).

**Figure 8.**
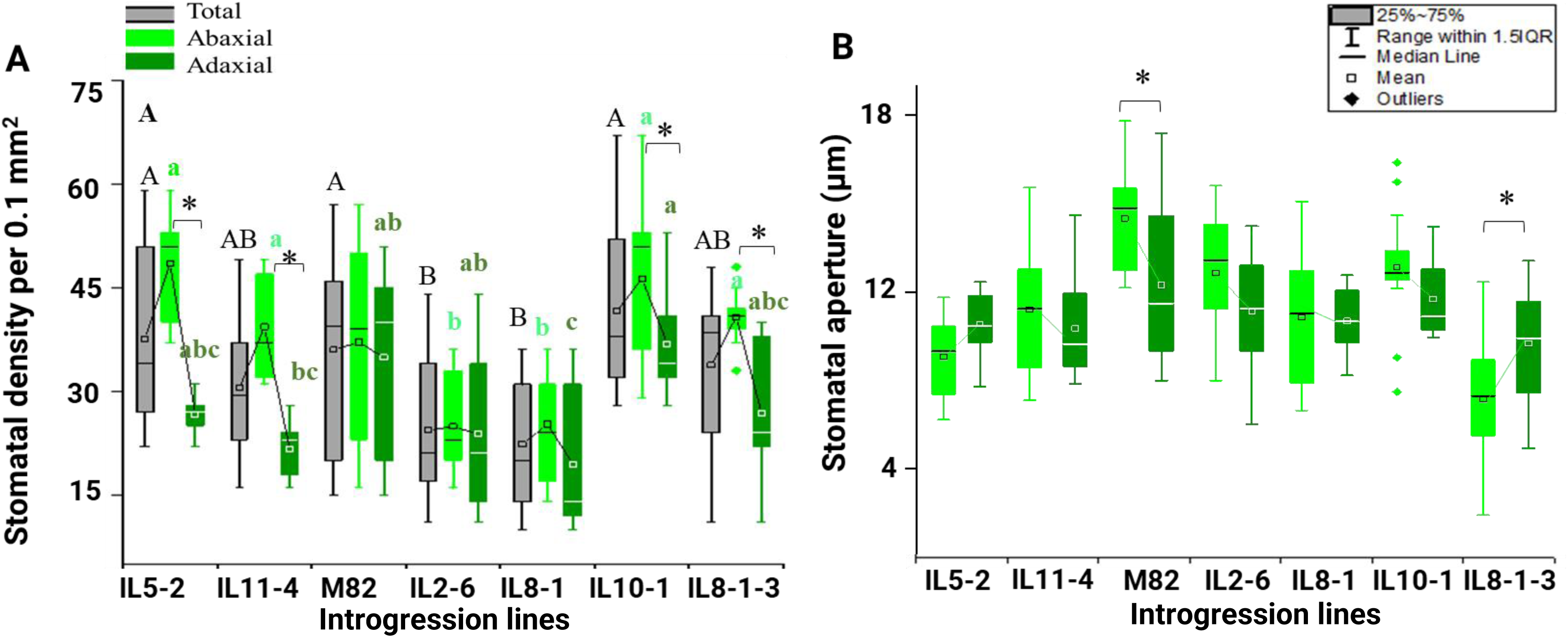
Leaf stomatal traits of the different tomato lines. A, Total (gray), abaxial (lower leaf side, light green) and adaxial (upper leaf side, dark green) stomatal densities of six introgression lines of tomato and M82. Data are derived from three technical and three biological replications imaged at their central lamina. The box-splitting horizontal bands indicate sample medians, the square box in the middle indicates the mean, and bars show the interquartile range (25th to 75th percentile), points below or above the interquartile ranges are outliers. B, Total stomatal apertures of the same lines (pore width, µm). * Indicates a significant difference between the abaxial and adaxial sides of an individual line, according to Student’s *t*-test. Different letters indicate significantly different means (capital letters for total stomatal density, and light green lower-case letters for the abaxial side of the leaves and dark green lower-case letters for the adaxial side), according to Tukey’s Honest Significance test (*p* < 0.05).

**Figure 9.**
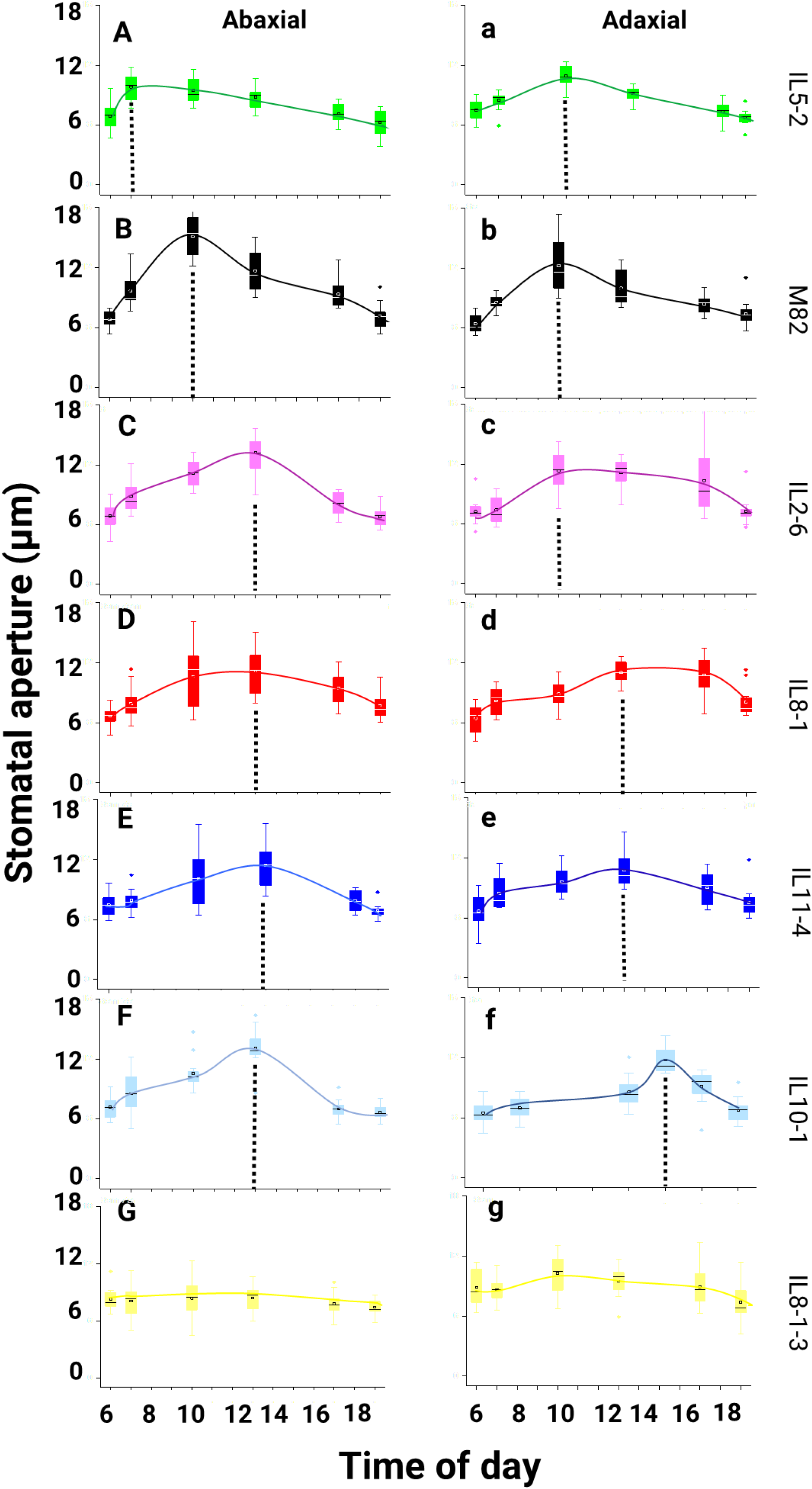
Daily variation in, A–G, abaxial stomatal apertures and, a–g, adaxial stomatal apertures. The broken lines indicate the maximum aperture during the daytime. The square (□) in the box plot represents the mean value. The box-splitting horizontal bands indicate sample medians and bars show the interquartile range (25th to 75th percentile). Points below or above the interquartile ranges are outliers. The dashed lines in each figure indicate the point on the day at which the apertures were largest.

### Abaxial, but not adaxial stomatal density, is correlated with the transcription of stomatal development candidate genes

We explored the relationship between transcriptome data generated by (Chitwood et al., 2013) and the abaxial and adaxial stomatal densities we measured. We discovered that the expression levels of the gene SPEECHLESS (SPCH) were strongly and positively correlated with abaxial stomata density (Figure 10 B and C). The Zeaxanthin epoxidase gene (ZEP) was strongly negatively correlated with abaxial stomatal density (Figure 10 E and F).

**Figure 10.**
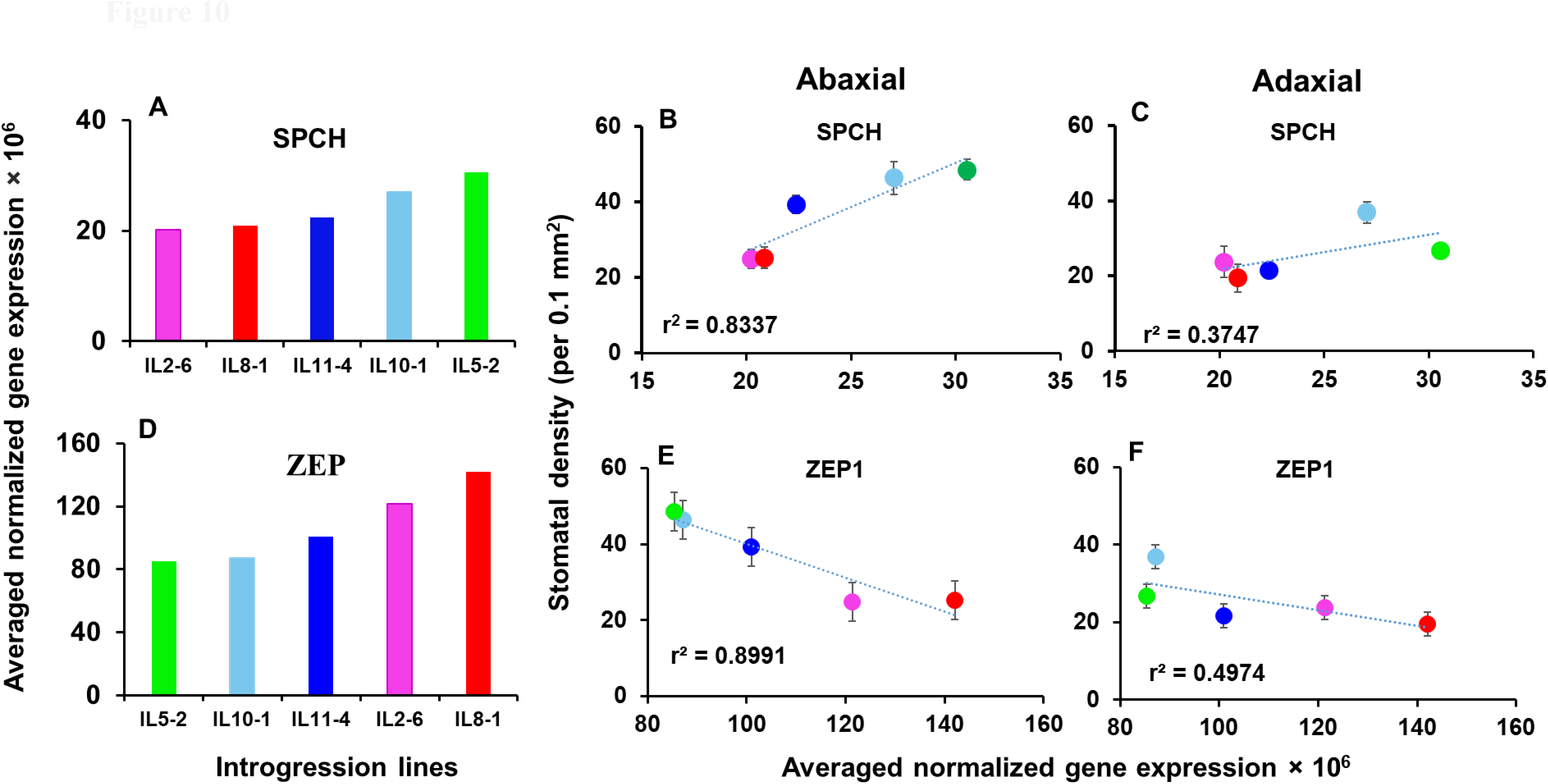
Expression of genes related to stomatal development correlated to abaxial and adaxial stomatal distribution. A, *SPCH* expression in the different lines. B, Relationship between *SPCH* expression and abaxial stomata density. C, Relationship between *SPCH* expression and adaxial stomatal density. D, *ZEP* expression in the different lines. E, Relationship between *ZEP* expression and abaxial stomatal density. F, Relationship between *ZEP* expression and adaxial stomatal density. The gene-expression data are from Chitwood et al. (2013).

## Discussion

Yield and biomass data, typically used to differentiate between genotypes grown in the field, are single endpoint traits. These data result from numerous G×E interactions throughout the growing season and can vary across locations and years (van Eeuwijk et al., 2019). This variability presents a significant challenge for crop-phenotype studies (Momen et al., 2019). To address some of these challenges, we selected seven different genotypes and analyzed at least four years of field data. The lines were categorized as high-yielders (IL5-2, IL11-4, and Il2-6), a medium-yielder (akin to M82), or low-yielders (IL8-1, IL10-1, and 8-1-3). They were continuously and simultaneously compared with each other and a control (i.e., M82) under identical atmospheric conditions in scenarios of optimal irrigation, drought stress, and recovery. To our knowledge, this is the first physiological phenotyping study incorporating multiple years of field performance across various genotypes.

Given that leaf photosynthesis is deemed crucial for yield improvement (Du et al., 2020; Tatsumi et al., 2020), we initially employed a manual gas-exchange apparatus to measure the A_N_ and gs of the selected lines (Supplemental Figure 1). While some trends were observed, no significant differences emerged that could elucidate the yield performance variations among these lines. This might be attributed to the limited sample size we could measure within the 2-hour window between 10:00–12:00, specifically only three biological replications for each of the seven genotypes. Due to the measurement’s nature, there was a ∼40 min gap between the three replications. Even within this brief period, environmental factors like sunlight levels, temperature, and VPD were not constant. Consequently, infrequent, manual measurements, even of a critical trait like AN, might not yield data suitable for robust selection criteria, especially when values differ in a naturally fluctuating environment (Sakoda et al., 2020).

The primary challenge of G×E phenotyping lies in detecting minor behavioral differences between lines under similar environmental conditions. Over time, these differences cumulatively impact crop yield (Moshelion, 2020; Gosa et al., 2022). Sampling longitudinal traits, i.e., traits measured repeatedly over time (Yang et al., 2006), is essential to discern how the environment influences plant performance. As reviewed by Moreira et al. (2020), measuring a crop’s longitudinal traits can better describe its dynamic nature. Consistent with this perspective, continuous measurements, as opposed to traditional single-point measurements, have been shown to enhance the accuracy of the longitudinal trait prediction model for plant shoot growth and aid in discovering loci associated with shoot growth trajectories (Campbell et al., 2019). To comprehend the dynamic trajectory responses of different lines, continuous monitoring of the soil-plant-atmosphere continuum (SPAC) conditions might be more informative than single time-point measurements. We observed that even under optimal irrigation, single-point measurements of daily transpiration could be misleading. Differences began to emerge over the growth period, suggesting that a single-point measurement during early growth might not offer much insight. This underscores the importance of identifying the appropriate time axis for evaluating any measurement data.

Another challenge with dynamic trajectory response is the need to compare different lines at similar drought stress levels, as plants exhibit varied water-consumption behaviors. Pursuing a deeper understanding of trait dynamics under water-deficit conditions is among the most significant challenges in breeding for higher yields under drought conditions (Sinclair, 2011; Snowdon et al., 2020). Designing a standardized, repeatable, and reliable drought experiment and screen remains a daunting task (Moshelion, 2020). In our study, we maintained similar drought treatments for all lines simultaneously (Figure 3B) by defining the initial drought-stress point in terms of each plant’s θcrit., as a standard drought evaluation point (Figure 4, Suppl. Figure 3). However, during the drought treatment, even though similar water content was automatically maintained for all lines (Figure 3 B), we observed a transpiration-positional inversion between the low- and high-transpiring plants (Figure 3 A, “transpiration flip-flop”).

This result highlights the challenges of the commonly used method of halting irrigation for several days to study drought stress, especially when involving plants of different sizes and environmental sensitivities. This outcome underscores the importance of caution when employing the method developed by Michael D. Snow and David T. Tingey (1985) to study water stress, especially when the experiment involves a variety of plant behaviors. Using θcrit. as a standard drought evaluation point (see Figure 4, Suppl. Figure 3) could help address this challenge. θcrit. is also crucial for demonstrating a plant’s dynamics and stress sensitivity, revealing how quickly high-transpiring plants can reduce their transpiration, potentially impacting their stress tolerance. For instance, as shown in Figure 4 and Suppl. Figure 3, IL5-2 and IL11-4 reached the θcrit. point more rapidly than other lines. This suggests that these lines are opportunistic, transpiring more under full irrigation and closing their stomata upon sensing a water deficit.

Under stress, plants typically transition from a productive to a survival mode, involving longitudinal physiological, anatomical, and biochemical adaptations (Kerchev and van Breusegem, 2021). Recovery from drought stress entails returning to pre-stress growth and physiological functioning levels once soil water content is restored. Although resilience is as vital as stress response, it has garnered less attention, possibly due to its characterization challenges (Guo et al., 2020).

As depicted in Figure 5 A, the examined lines displayed varied abilities to revert to their pre-stress daily transpiration levels compared to their daily transpiration during the first five days post-irrigation resumption. Lines with higher pre-stress transpiration levels (IL5-2, IL11-4) recovered swiftly post-drought, contradicting our hypothesis that higher-transpiring plants would be more vulnerable to drought and recover more slowly. Similarly, these lines were classified as highly resilient and medium resilient in the field (Table 1) and exhibited faster stomatal closure, higher θcrit. (Figure 4, Suppl. Figure 3), higher WUEb (Figure 5c), and relatively high Gsc to TR under drought conditions (Figure 7 A, a and B, b, respectively). This indicates the importance of a rapid and dynamic response to water stress.

Based on these findings, we infer that stress-adaptation mechanisms, which might be elusive during the stress period, could contribute to plant resilience. This aligns with a previous report on maize (Chen et al., 2016), which posited that drought recovery is integral to whole-plant growth under water-stress conditions. Employing a method to detect drought-stress resilience during early growth stages, combined with field trials, might enable resilience prediction during earlier growth stages. This approach could also help identify genetic variability in novel tolerance mechanisms. If paired with field testing (Chapuis et al., 2012), data should be gathered during both the vegetative and reproductive plant growth phases, as plants tend to exhibit varied stress responses during these distinct phases (Gosa et al., 2022; Chen et al., 2016).

### Dynamic Water-Use Efficiency: The Morning Stomatal Peak Under Water-Deficit Conditions

Plants demonstrate dynamic water-use regulation throughout the growing season in response to VPD changes. The temporal dynamics of water-use traits can significantly enhance productivity (Sinclair, 2018). Similar to seasonal variation, diurnal VPD is dynamic, influencing instantaneous plant responses (Figure 6). However, in the absence of SWC limitations, all lines showed analogous response patterns, differing only in their absolute values (Figure 6 B, C). Under limiting SWC, different lines displayed varied responses to VPD and PAR (Figure 7 and Supplemental Figure 2), suggesting that certain dynamic responses might be more advantageous in specific environments. A notable example of such beneficial dynamic responses is observed in the early morning whole-canopy conductance peaks of the high-yielding lines. Under drought conditions, IL5-2 and IL11-4 (Figure 7 A and B) maintained relatively high whole-canopy conductance when PAR was high, but VPD was still low. This resulted in a relatively low rate of water loss to transpiration (Figure 7 a and b). A morning peak of this nature was reported in solanum penelli (Lupo and Moshelion, 2023).

This plastic trait is likely to have been reintroduced through introgression. In other words, it’s probable that introgression led to the recovery of this trait, and the diversity lost during the genetic bottlenecks that occurred since the domestication of the tomato, as suggested by Gramazio et al., 2021. Such opportunistic stomatal behavior has been documented in highly water stress-tolerant forest plants like Acacia and hemiparasitic mistletoes (Loranthus europaeus (LE)) (Ullmann et al., 1985; Resco de Dios et al., 2016), a high-yielding wheat (Triticum durum cv) introgression line (Bacher et al., 2021), soybean (Glycine max) (TEARE and KANEMASU, 1972), and Arabidopsis thaliana (Hassidim et al., 2017). While these studies acknowledged the existence of this stomatal morning-rise phenotype, it was hypothesized that this phenotype would boost productivity and fitness under drought by utilizing the photosynthetically active radiation (PAR) to enhance CO2 assimilation (Schoppach et al., 2020). In a recent study, we demonstrated that this type of early morning Gsc peak, termed the “golden hour” (Gosa et al., 2019), is strongly correlated with tomato yield in the field (Gosa et al., 2022).

Our current findings underscore the significance of this early morning stomatal peak and highlight genetic variability leading to this crucial trait. Specifically, the two high-yielding lines were IL5-2 and IL11-4, which are otherwise distinct from each other. Moreover, these lines also displayed high WUEb, high shoot dry weights (Figure 5B), and medium to high drought tolerance in the field (Table 1). In contrast, another high-yielding line, IL2-6, presented daily Gsc–Tr kinetics that were similar to the low-yielding Il8-1 (Figure 7). IL2-6 also had low WUEb and low shoot dry weights (Figure 5B), as well as low drought tolerance in the field (Table 1). Thus, different combinations of various traits resulted in similar outcomes.

In recent years, several studies have proposed stomatal functional anatomy mechanisms as promising targets for enhancing productivity and resilience (Sakoda et al., 2020b; Sultana et al., 2021). For instance, high stomatal density has been suggested as a safety-efficiency trade-off, as it exhibits greater sensitivity to closure during leaf dehydration (Henry et al., 2019). Another mechanism proposed as a good proxy for productivity and WUE is the ratio between the stomatal densities on the abaxial and adaxial sides of a plant’s leaves (Muir et al., 2014).

Indeed, our results revealed different combinations of both in the different IL lines. All the IL lines tested had similar total stomatal densities. However, higher stomatal densities on the abaxial side were observed among both high-yielding lines (L5-2 and IL11-4) and low-yielding lines (IL10-1 and IL 8-1-3; Figure 8 A). Additionally, other high-yielding lines, M82 and IL2-6, had stomatal densities and stomatal ratios similar to the low-yielding IL8-1 (Figure 8 A).

These findings prompt further questions about the roles of stomatal ratio and stomatal density in Gsc regulation. Although the abaxial stomata seem to play pivotal roles in light sensitivity, photosynthesis, and WUE (TURNER, 1970; Driscoll et al., 2006; Wang et al., 2008; Lei et al., 2018), the actual relative contribution of the abaxial and adaxial leaf sides to crop productivity remains ambiguous. It’s influenced by several factors, such as the position of light illumination in the greenhouse, wind movement in the field, and crop type (Zhang et al., 2016; Paradiso et al., 2020). We propose that, in tomatoes, the combination of stomatal ratio, stomatal density, and the sensitivity of the stomatal conductance on each leaf side to ambient conditions is key to understanding a genotype’s adaptation to a specific environment (G×E optimization), its resilience, and its WUE. For instance, both IL8-1 and IL8-1-3 were very low in all productive and resilient field-based parameters (Table 1). Indeed, their daily transpiration levels were low, and despite their shorter exposure to stress, they were the last to recover (Figure 3). Both lines exhibited similar low shoot dry biomass and low WUEb (Figure 5). Moreover, under stress, both displayed similar patterns of daily Gsc (Figure 7 E and G) and Tr rates (Figure 7 e and g). However, these similar response patterns result from different stomatal ratios and stomatal densities (Figure 8 A) and maximal stomatal apertures at different times of the day (Figures 8 B and 9 D, G, d, and g). Specifically, IL8-1 had a lower stomatal density but larger stomatal apertures on the adaxial sides of its leaves, which have more exposure to the atmosphere during the highest VPD hours. It maintained those large apertures for a longer time (Figure 9 d), resulting in a transpiration rate similar to IL8-1-3 (Figure 7 e and g), which had a higher stomatal density and stomatal ratio (Figure 8), yet lower daily aperture kinetics on both the abaxial and adaxial sides of its leaves (Figure 9 G and g).

On the other hand, IL5-2, which we categorized as the idiotype (high levels of all productivity and resilience parameters; Table 1), exhibited. high levels of daily transpiration, even though it was exposed to drought stress for a longer period (earlier θcrit.; Figure 4, Suppl. Figure 3). Plants of this line also recovered the quickest (Figure 3 A) and had high levels of dry biomass and WUEb (Figure 5 C). Accordingly, under stress, this line exhibited a high Gsc but low Tr rates (Figure 7 A and a) due to an earlier aperture peak (Figure 9 A). Line IL11-4 was almost identical to IL5-2 in all parameters, including a similar stomatal ratio and stomatal density (Figure 8 A), yet it had a later aperture peak (Figure 9 B and b) at a higher VPD. This “risk-taking” may reflect its medium shoot weight, tolerance, and resistance in the field.

The other lines (i.e., M82 and IL2-6) were similar in their daily transpiration, dry biomass, WUEb, θcrit., daily Gsc and TR patterns, and stomatal ratios (Figures 3, 4, 5, 7, and 8, respectively). However, IL2-6 had a lower stomatal density (Figure 8) and larger adaxial stomatal apertures late in the afternoon, which could explain why it was slower to recover (Figure 3 A) and could be related to its medium level of tolerance under field conditions. IL10-1 was also like M82 in its daily transpiration, dry biomass, WUEb, θcrit., stomatal density, and adaxial–abaxial stomatal aperture ratio (Figures 3, 4, 5, 8, and 9, respectively), yet IL10-1 had a higher stomatal ratio (Figures 8) and much higher daily Gsc and TR under drought conditions (Figure 7 F and f). These patterns could be related to its low tolerance in the field (Table 1). IL8-1-3 had a stomatal density and stomatal ratio like the high-yielding IL5-2 (Figure 8 A), but it exhibited low transpiration (Figure 3 A) and low yield in the field (Table 1). This could be explained by the fact that this line had relatively small stomatal apertures on both sides of its leaves (Figures 8 B and 9 G and g). Collectively, these findings suggest that the combination of stomatal density, stomatal ratio, daily stomatal-aperture profile, and maximal stomatal apertures is essential for plant adaptation and productivity under water-stress conditions.

To shed light on the potential genetic basis of this stomatal trait variation as a target for future study, we utilized transcriptome data for these tomato lines generated by (Chitwood et al., 2013). We observed a correlation between two of the stomatal development-related genes and abaxial and adaxial stomatal density. These genes are SPEECHLESS (SPCH Solyc03g007410.2.1) and a gene that encodes zeaxanthin epoxidase (ZEP, Solyc02g090890.2.1).

SPCH is a signal integrator, which combines intralineage signals with hormones and environmental effects during development to regulate stomatal density and patterning (Chen et al., 2020). Zeaxanthin epoxidase (ZEP) is involved in the biosynthesis of abscisic acid and the xanthophyll cycle, playing a crucial role in regulating plant responses to various environmental stresses (Lou et al., 2017). Interestingly, we observed a strong positive relationship between abaxial stomatal density and the expression of the stomatal developmental gene, SPCH. In contrast, the correlation between adaxial stomatal density and SPCH expression was less pronounced (Figure 10 C and D). Additionally, we noted a strongly negative correlation between abaxial stomatal density and ZEP expression. ZEP was expressed at a very low level in IL5-2, the most plastic line in this study (Figure 10 D–F). Although our current work did not delve into the elucidation of genetic mechanisms, we believe that these observed correlations will stimulate future research, encouraging researchers to explore this subject further. Understanding how stomatal ratios can be used to enhance crop plasticity in a constantly changing environment is of paramount importance.

In conclusion, plants with high levels of transpiration can be more productive in the absence of water limitations. However, the key to maximizing productivity under SWC constraints is to maximize stomatal aperture when VPD is low, quickly close stomata when VPD increases and/or SWC decreases, and then rapidly open stomata when SWC increases. Yet, continuous measurement of stomatal apertures is not feasible as part of a phenotypic scan. Similarly, measuring stomatal conductance with a manual, steady-state, gas-exchange apparatus is also not feasible due to the limited areas these machines measure (typically a leaf), which doesn’t reflect the functional anatomy of the entire plant (including the abaxial-adaxial stomatal density ratio, leaf structure, canopy phyllotaxis, and other factors). Conversely, whole-plant measurement by the gravimetric method provides an absolute measurement of the whole plant’s response to environmental changes. This can be invaluable in high-throughput screening for either forward or reverse G×E phenotyping, greatly benefiting breeding programs aimed at improving crop performance in diverse climates.

## Acknowledgements

This research was supported by the ISF-NSFC joint research program (grant no. 2436/18), as well as the Israel Science Foundation (grant no. 1043/20) and the Israel Ministry of Agriculture and Rural Development (Eugene Kandel Knowledge Centers) as part of the Root of the Matter: The Root Zone Knowledge Center for Leveraging Modern Agriculture. We thank Prof. Dani Zamir at our institute for providing us with the IL seeds and multiple years of field-based yield data for the lines used in this study.

## Author Contributions

Sanbon Chaka Gosa planned, conducted, and analyzed all the experiments, wrote this thesis except the following:

Bogale Abebe and Ravitesjes Patil took part in conducting the lysimeter experiments. Ramon Manica took part in stomatal imprint and leaf gas exchange experiments.

Moshelion M: corresponding author, supervised, planned wrote the MS together with Sanbon Gosa.

## Supplementary

**Supplementary Figure 1.**
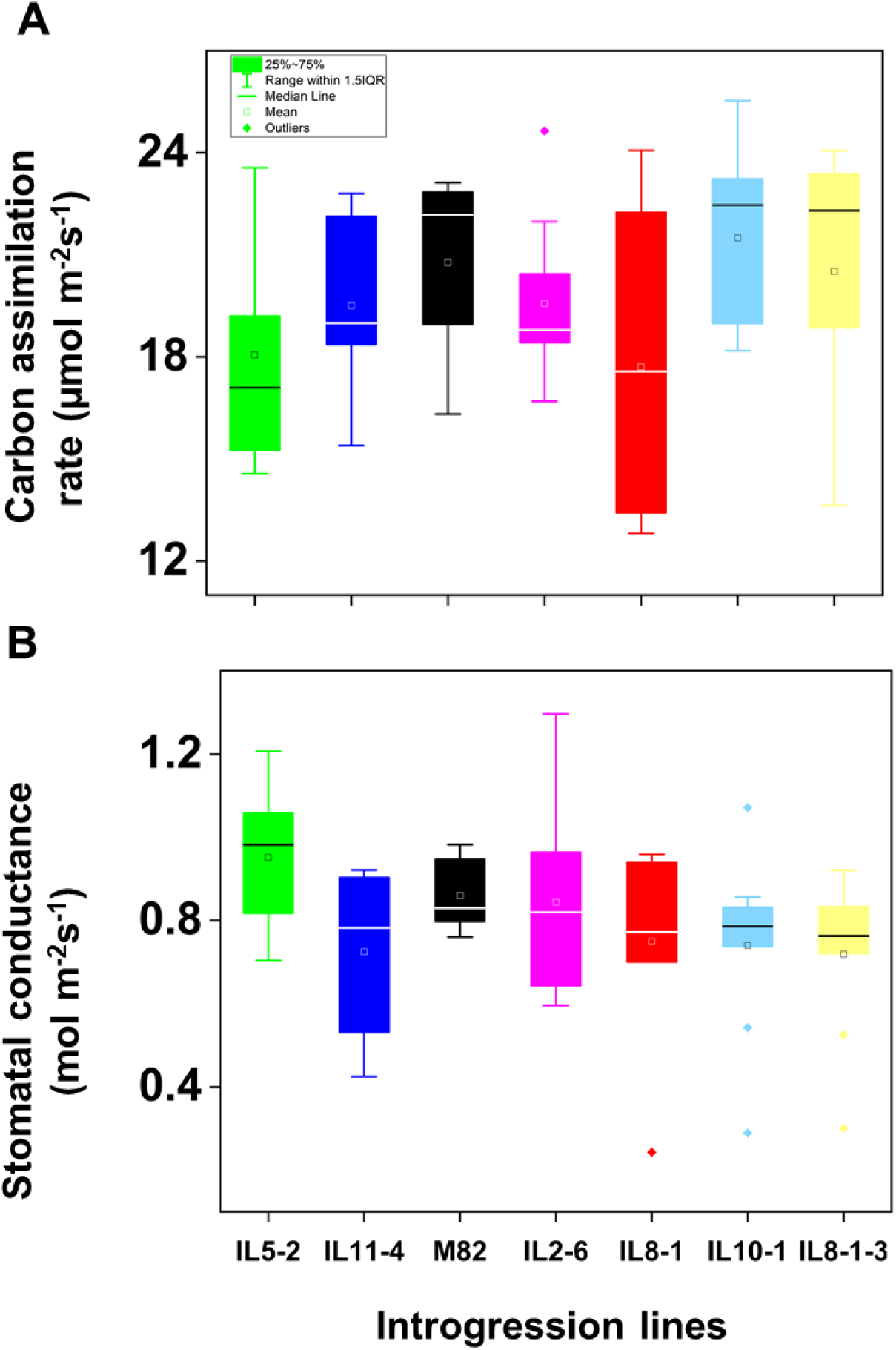
Leaf gas exchange of different tomato introgression lines under well-irrigated greenhouse conditions. A, Rate of carbon assimilation in young leaves before flowering. B, Stomatal conductance of young leaves before flowering. The square (□) in the box plot represents the mean value. The box-splitting horizontal bands indicate sample medians and bars show the interquartile range (25th to 75th percentile), points below or above the interquartile ranges are outliers.

**Supplementary Figure 2.**
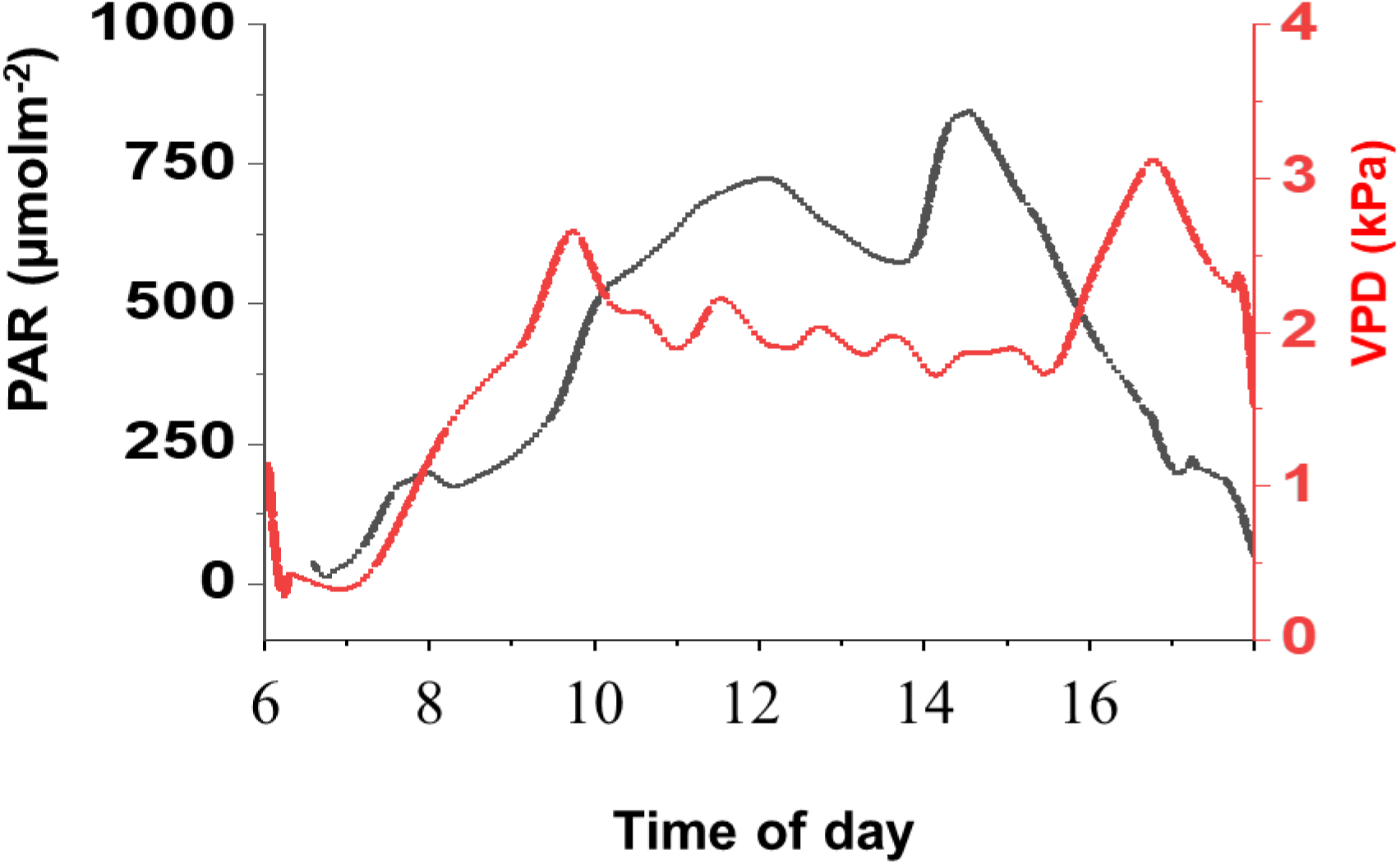
Daily patterns PAR (black line) and VPD (red line) during water stress period.

**Supplementary Figure 3.** The physiological drought point (θcrit.) of individual introgression lines, calculated by piecewise correlation fit between volumetric water content (VWC) and transpiration rate using ‘SPAC Analytics’ software (for all of the plants presented in Fig. 4).

