## Supplemental Figure 1 for "Diurnal stomatal apertures profile and density ratios affect whole-canopy conductance, drought response, water-use efficiency and yield"

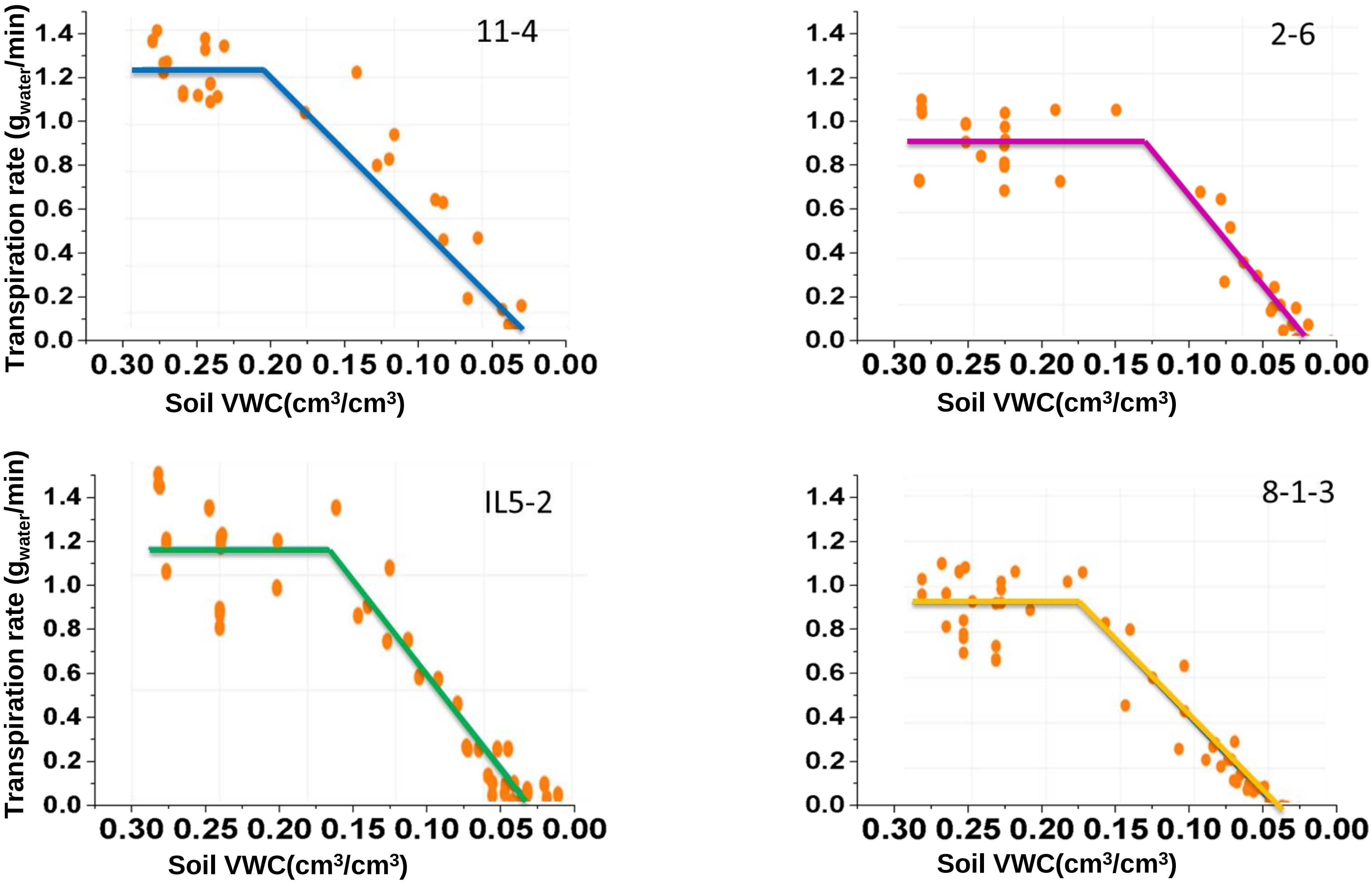


**Supplementary Figure 3.** Theta crit point of individual introgression lines, calculated by piecewise correlation fit between volumetric water content (VWC) and transpiration rate using 'SPAC Analytics' software.

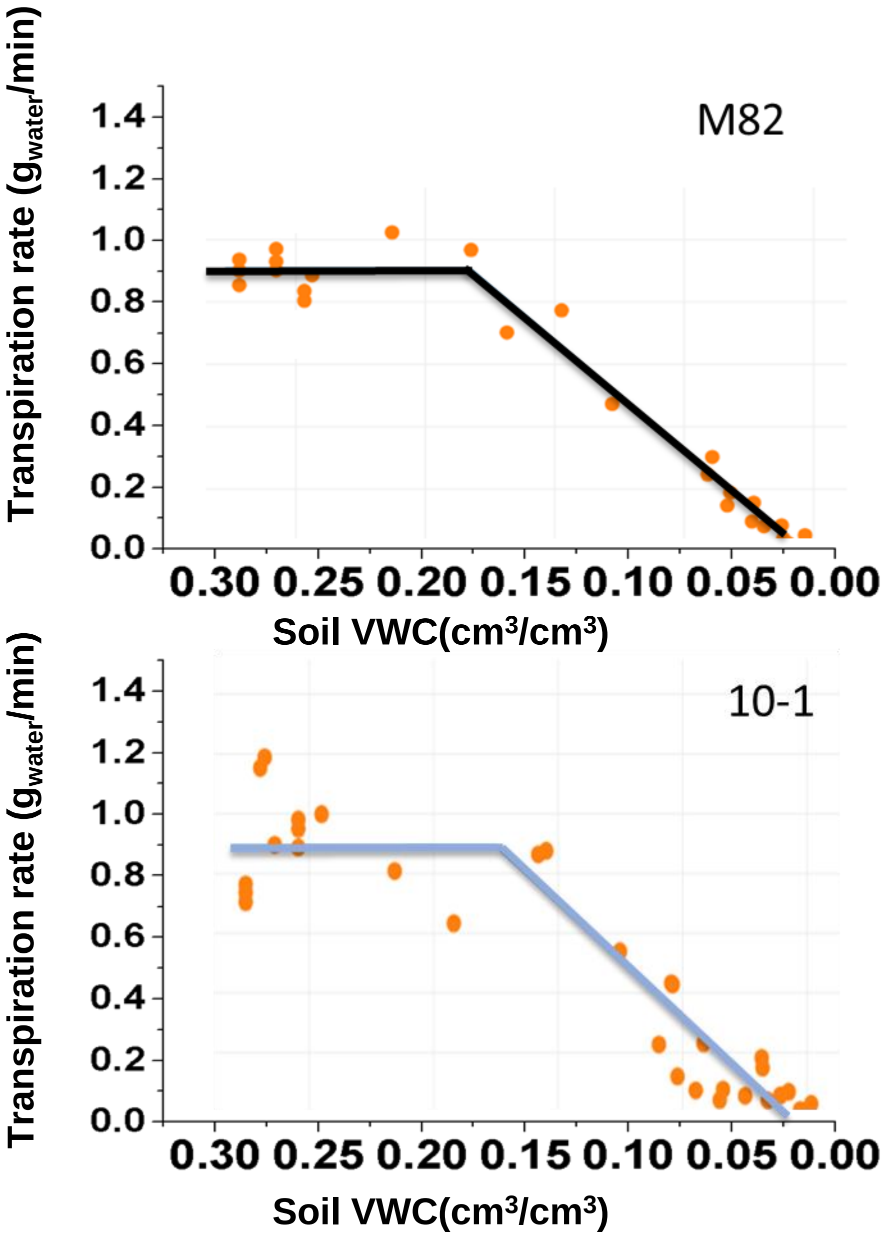


**Supplementary Figure 3.** Theta crit point of individual introgression lines, calculated by piecewise correlation fit between volumetric water content (VWC) and transpiration rate using ‘SPAC Analytics’ software.
