## Supplemental Materials and Methods for "Diurnal stomatal apertures profile and density ratios affect whole-canopy conductance, drought response, water-use efficiency and yield"

### Field experiments

In this study, we utilized 30 tomato genotypes: 29 ILs derived from crossings of Solanum pennellii and the cultivar M82 (Solanum lycopersicum cv. M82; Table 1). Each line contained a single homozygous restriction fragment-length polymorphism of the Solanum pennellii chromosome segment (Eshed and Zamir, 1995). For the genotype-performance screening of seven selected genotypes, we employed historical yield data from field experiments conducted in 1993, 2000, 2001, 2002, 2003, 2004, and 2010. Briefly, the open-field experiments took place at the Western Galilee Experimental Station in Akko, Israel, during the summer, following a randomized block design (Gur and Zamir, 2004). These trials maintained a planting density of one plant per $m^2$. Both wet and dry fields began the growing season at field water capacity. As there was no rainfall during the experimental period, only the irrigation system managed the drought scenarios. Details of the agricultural practices and measurements have been previously published (Fridman et al., 2000; Gur and Zamir, 2004).

### Leaf gas-exchange measurements

Plants were cultivated in the ground (sandy-loam soil) within a semi-controlled greenhouse at the Hebrew University of Jerusalem, Faculty of Agriculture, Rehovot, Israel. The genotypes were planted and measured in random order. We assessed leaf gas-exchange on the youngest, fully extended leaf. Data were collected from mature, fully expanded leaflets at the canopy top of ~8-week-old plants between 10:00 am and 12:00 pm. A portable infrared gas analyzer (LI-6800XT; Li-Cor Inc., Lincoln, NE, USA) was employed to determine the carbon assimilation rate ($A_N$). Stomatal conductance (gs) was measured in a 6-$cm^2$ chamber at midday, with the $CO_2$ reference set at 400 mmol $m^{-2}$ $s^{-1}$, PAR at 400 mmol $m^{-2}$ $s^{-1}$, VPD at 1.4 kPa, and temperature at 25°C. These conditions were chosen to replicate the greenhouse's environmental conditions during the initial measurements.

### Reverse phenomics using the physiological-phenotyping platform in a greenhouse

### Experimental setup

Five-week-old seedlings of eight selected ILs and M82 tomato plants were transplanted into pots and grown in a greenhouse affiliated with the Israeli Center of Research Excellence (ICORE) for Plant Adaptation to the Changing Environment. This was located at The Hebrew University of Jerusalem, Faculty of Agriculture, Rehovot, Israel, during September 2019. An overview of the nutrients supplied to the plants via the irrigation system (fertigation) is presented in (Dalal et al., 2020). Before the experiment's commencement, all load-cell units were calibrated for accuracy and drift level under constant load weights (1 kg and 5 kg) using the Plantarray auto-calibration application. Atmospheric conditions and the soil were monitored (see Fig. 1), as described in (Dalal et al., 2020).

The setup included highly sensitive, temperature-compensated load cells, used as weighing lysimeters. Each controller was connected to its control unit, which collected data and managed irrigation. A 4-L pot containing a single plant in 20/30 sand (Negev Industrial

Minerals Ltd., Israel) as a growth medium was placed on each load cell. The numbers 20/30 refer to the upper and lower size of the mesh screen through which the sand passed (20 = 20 squares across one linear inch of screen), resulting in a sand particle size between 0.595- and 0.841-mm. Fertilizer (poly feed 17:10:27, Haifa Chemicals, Haifa, Israel) was provided to the plants through the irrigation system (fertigation). The containers, which fit the pots tightly to prevent evaporation, had orifices at different heights on their side walls to allow for different water levels after the drainage of excess water post-irrigation. Evaporation from the pot surface was prevented by a cover with a circle cut out at its center, allowing the plant to grow. All pots were fertigated by four drippers, which were inserted into the upper part of the sand to ensure even wetting during each irrigation event. Fertigation occurred during the night in multi-pulses (i.e., the fertigation in the control treatment consisted of four irrigation pulses for 15 min, every 2 h to ensure proper leaching and reaching full pot water capacity).

**Drought treatment**

Since each individual plant had a unique transpiration rate based on its size and location in the greenhouse, halting the irrigation for all plants simultaneously would result in a non-uniform drought treatment. To ensure a standardized drought treatment (i.e., similar drying rate for all pots), drought scenarios were automatically controlled via the system's feedback-irrigation controller. For the drought treatment, the system was set to irrigate each plant to 80% of its previous day's transpiration, ensuring all plants experienced the same gradual water stress [see Figure 3B in (Dalal et al., 2020)].

**Measurement of quantitative physiological traits**

The plant water-relations kinetics (recorded by the system every 3 min) and quantitative physiological traits of the plants were determined simultaneously for all plants (Figure 1), following (Halperin et al., 2017) with minor modifications. The examined traits included daily transpiration, transpiration rate (TR), whole-canopy stomatal conductance (Gsc), and biomass water-use efficiency (WUEb). Cumulative transpiration (CT) was calculated as the sum of daily transpiration for all the experiment days for each plant.

Biomass water-use efficiency (WUEb) was determined by dividing each plant's dry biomass weight by its CT at the experiment's conclusion, as defined in (Leakey et al., 2019). Daily weight gain was calculated by subtracting each day's pot weight from the pot weight measured on the previous day, both after reaching field capacity and full drainage at 04:00 [as detailed in (Halperin et al., 2017)]. The plant's recovery from drought was described by the recovery of daily transpiration to its pre-drought level after irrigation resumed. The recovery rate was determined by comparing the amount of daily transpiration for 5 days post-recovery.

Calculated volumetric water content (Cal. VWC) was derived from the mass balance difference of reserve water, soil wet weight, soil dry weight, and soil volume (see Dalal et al., 2020). Theta crit. (θ) is the point at which transpiration begins to be affected by limited

soil water availability. It was determined by the piecewise linear fit of the transpiration rate and calculated VWC of the plants subjected to the drought treatment.

**Stomatal density and aperture**

A rapid imprinting method (Geisler and Sack, 2002) was employed to determine stomatal apertures and density (Figures 8 and 9). Briefly, light-bodied vinyl polysiloxane dental resin (Heraeus-Kulzer; Hanau, Germany) was attached to the abaxial and adaxial sides of leaves and then removed after drying for 1 min. Mirror images of the resin imprints were created using nail polish. Once dried, the nail polish was lifted from the resin epidermal imprints. The nail-polish imprints were mounted on microscope slides. A Leica DM500 microscope equipped with a 40x objective and a Leica ICC50W camera was used to observe and photograph the imprints at 20x and 60x magnification for stomatal density and apertures, respectively. The stomata in a field of view were counted at 20x magnification. Using the ImageJ software (http://rsb.info.nih.gov/ij/), stomatal images were analyzed to determine aperture size. A microscopic ruler (Olympus; Tokyo, Japan) was used for size calibration.

**Statistical analysis**

Continuous data were filtered and summarized using the SPAC analytic software embedded in the Plantarray system (PlantDitech, Yavne, Israel). All analyses were conducted using the JMP® 15.0 Pro statistical package (SAS Institute, Cary, NC, USA) unless otherwise specified. Box plots and continuous line graphs were generated using OriginPro, Version 2021 (OriginLab Corporation, Northampton, MA, USA).
